# Persistent Activation of Endothelial Cells is Linked to Thrombosis and Inflammation in Cerebral Cavernous Malformation Disease

**DOI:** 10.1101/2025.06.29.662238

**Authors:** Helios Gallego-Gutierrez, Eduardo Frias-Anaya, Cassandra Bui, Louie Zhao, Emily Hsu, Hannah S. Indralingam, Jakob Körbelin, JoAnn Trejo, Jeffrey Steinberg, Nathan R. Zemke, Miguel A. Lopez-Ramirez

## Abstract

**BACKGROUND:** Cerebral cavernous malformations (CCM) are neurovascular lesions that affect both children and adults, and morbidity often results from thrombosis, bleeding, and neurological dysfunction. Studies indicate that inflammation-related activation of endothelial cells contributes significantly to the worsening of CCM disease. This suggests that ongoing vascular inflammation and endothelial dysfunction are key factors associated with thrombosis and bleeding in CCM disease. However, the inflammatory mechanisms leading to altered brain endothelial cell function with a high propensity for thrombosis, inflammation, and dysfunction are not fully understood.

**METHODS:** Multi-omic analyses was conducted by performing simultaneous high-throughput single-nucleus RNA sequencing (snRNA-seq) and single-nucleus transposase-accessible chromatin sequencing (snATAC-seq) with the 10x Genomics multiome platform in combination with immunofluorescence to study CCM pathogenesis in both female and male mice with CCM (*Slco1c1-CreERT2; Pdcd10^fl/fl^*) disease. The analysis was complemented with bulk RNA-seq, bulk ATAC-seq, and ChIP-seq (Chromatin immunoprecipitation sequencing) using an in vitro human CCM model. An AAV-BR1 viral system selectively upregulates the activator protein-1 (AP-1) transcription factor JUNB in brain endothelial cells was used to evaluate its effectiveness in maintaining a persistent activated cell state during the pathogenesis of CCM.

**RESULTS:** We found that epigenetics significantly influences the subtype identity and function of brain endothelial cells within the arteriovenous axis. Through multi-omic analyses, specific regulatory elements and enhancers (cis-Regulatory Elements, cCREs) in mouse brain endothelial cells were identified that influence subtype-specific transcriptional programs and the transcription factors responsible for establishing the various subtypes of brain endothelial cells. Additionally, large-scale epigenomic reprogramming of brain endothelial cell subtypes was observed during the pathogenesis of CCM disease. Among the most significant changes were alterations in the chromatin state of endothelial cells, along with transcriptional processes associated with a persistently activated endothelial cell state, which renders them susceptible to inflammation and thrombosis. The activator AP-1 transcription factor JUNB was identified as a key regulator of the persistently activated endothelial state during chronic neuroinflammation. Moreover, both trans- and cis-regulatory factors conserved between mice and humans were discovered and contribute to the progression of chronic CCM disease.

**CONCLUSIONS:** Epigenetics plays a crucial role in determining the transcription patterns and functions of brain arteriovenous endothelial cells. The activator JUNB is identified as a driver of chronic brain vascular inflammation by inducing a persistent activated endothelial cell state from epigenome reprogramming.

**Graphical Abstract:** 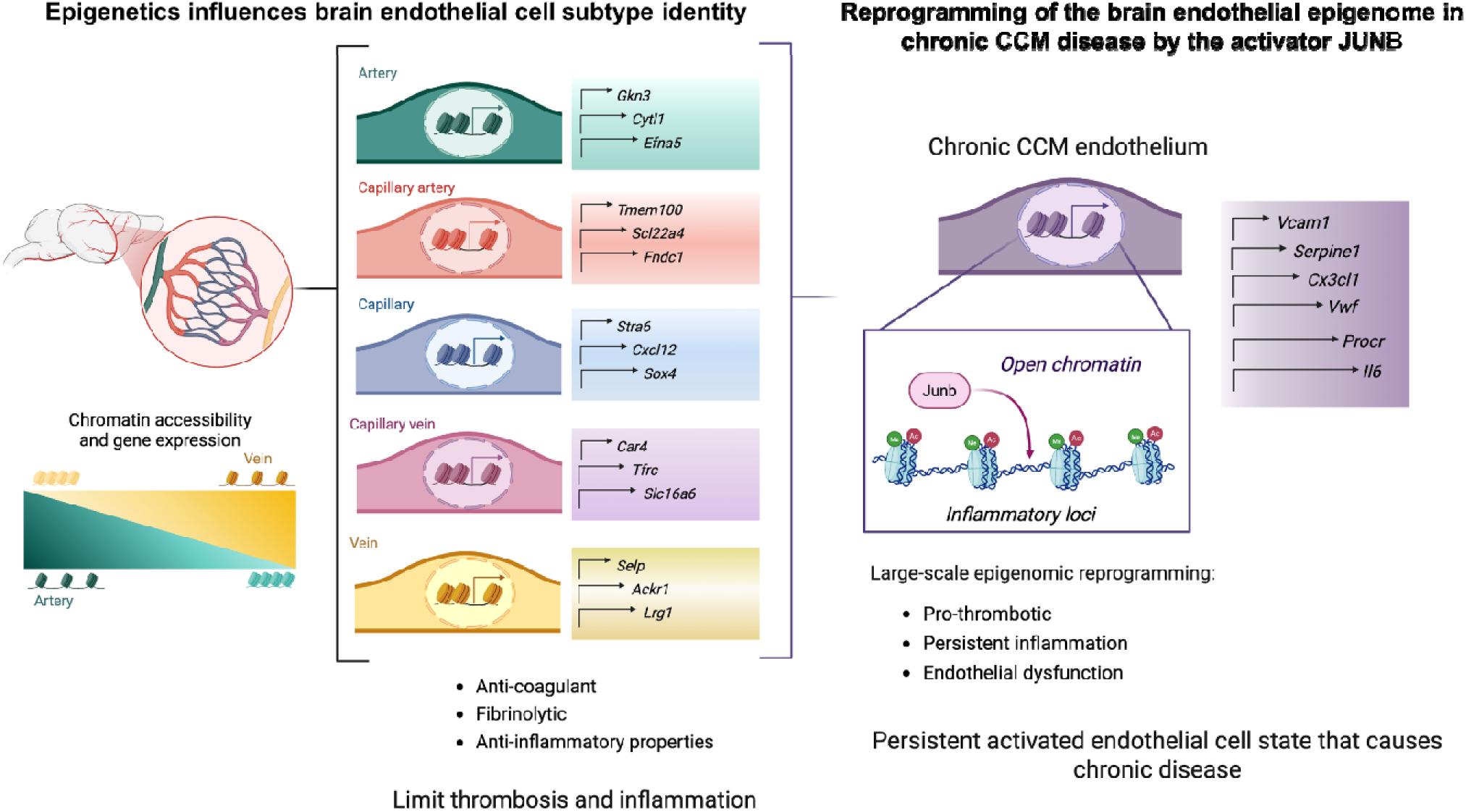

## Introduction

Cerebral cavernous malformations (CCMs) are common neurovascular lesions associated with lifelong risk of brain hemorrhage, thrombosis, and neurological deficits. An estimated 0.5% of the US population develops a CCM lesion, but only 25% are diagnosed(*1r-3*), with approximately 20% of cases identified during childhood(*2*). Familial CCM is an autosomal dominant disease with variable penetrance, even within the same family(*4*). Familial cases typically exhibit multiple lesions, whereas sporadic cases (approximately 80% of CCM cases) usually present with an isolated lesion. The heterogeneity in disease severity suggests that environmental, genetic polymorphisms, and biological factors act as disease modifiers(*3, 5-10*). The most common symptoms observed in patients with symptomatic CCMs (familial or sporadic) include motor disability, weakness, seizures, stress, and anxiety(*3, 11*). Currently, there is no effective pharmacological therapy for preventing or treating CCM, and surgery is the only treatment option for some patients(*5, 12, 13*).

The vascular hypothesis establishes that CCMs triggered by loss-of-function mutations (which can be inherited as germline or somatic mutations) in the genes *KRIT1* (Krev1 interaction trapped gene 1, *CCM1*), *CCM2* (*Malcavernin*), or *PDCD10* (Programmed cell death protein 10, *CCM3*)(*14-17*) propel brain vascular changes (CCM endothelium), characterized by disrupted intercellular junctions(*18-21*), increased angiogenesis(*19, 22-25*), endothelial-mesenchymal transition(*26*), reactive oxygen species (ROS)(*14, 27-29*), and vascular permeability(*18, 30-41*). These processes can lead to hemorrhage(*42, 43*) and inflammation, which are key characteristics of CCM lesions observed in both human cases and CCM animal models(*12, 17-19, 44-46*). Studies have highlighted inflammation as a key contributor to the development of chronic CCM disease(*45, 47-52*) and morbidity, typically attributed to bleeding, inflammation, and thrombosis(*45, 47, 48, 53*). This raises the possibility that inflammation-induced vascular changes may significantly contribute to the chronic nature and progression of CCM disease, and that persistent endothelial cell activation may be crucial in maintaining this chronic state.

Additionally, various signaling pathways, transcription factors, and likely alterations in chromatin accessibility may regulate the complex transcriptional changes in CCM endothelium(*29, 51, 54-56*) that contribute to an activated endothelial cell state. Recently, epigenetic changes have also emerged as important regulators of chronic inflammation(*57*) and are associated with the development of brain vascular malformation(*56, 58*). Targeting epigenetic changes has been proposed as a novel therapeutic approach in CCM disease(*56, 59*).

The Activator Protein-1 (AP-1) family consists of heterodimers and homodimers formed by basic region leucine zipper protein interactions. The most common combinations include Jun-Jun, Jun-Fos, and Jun-ATF dimers(*60*). These transcription factors bind to the promoters of target genes in a sequence-specific manner, allowing them to modulate gene expression and promote chromatin remodeling(*61-64*) through direct interaction with chromatin remodelers(*65-70*). AP-1 transcription factors are regulated by external signals and can act as activators in cis-regulatory regions to promote Th17 cell differentiation in chronic neuroinflammatory diseases(*71, 72*). Among the AP-1 transcription factors, JUNB family member has been shown to contribute to persistent inflammation(*73-76*). *Junb* is upregulated following intracerebral hemorrhage (ICH) in animal models, and this upregulation is linked to long-term neurological deficits caused by ICH(*77*). However, its specific role in the persistent activation of endothelial cells remains unclear.

Here, we report that epigenetics plays a significant role in influencing the identity and function of brain endothelial subtypes within the arteriovenous axis. Through multiomics analyses, we observed extensive epigenomic reprogramming of brain endothelial cell subtypes in a chronic model of CCM disease, characterized by an extensive neuroinflammatory profile. We identified the brain endothelial AP-1 transcription factor JUNB as an activator that regulates inflammatory loci and maintains a persistent activated endothelial cell state in chronic neuroinflammatory CCM animal models. Additionally, we uncovered both trans- and cis-regulatory factors that are conserved between mice and humans, contributing to the progression of chronic CCM disease and susceptibility to inflammation and thrombosis.

## Material and Methods

### Genetically modified animals

Brain endothelial-specific conditional *Pdcd10*-null mice were generated by crossing a Slco1c1 promoter-driven tamoxifen-regulated Cre recombinase (*Slco1c1-CreERT2*, a gift from Markus Schwaninger, University of Lu beck) strain with loxP-flanked Pdcd10 (*Pdcd10*^fl/fl^, a gift from Wang Min, Yale University; *Slco1c1-CreERT2;Pdcd10*^fl/fl^) mice. Brain endothelial-specific conditional tdtTomato reporter mice were generated by crossing B6.Cg-Gt(ROSA)26Sor^tm14(CAG-tdTomato)Hze^/J and *Slco1c1-CreERT2;Pdcd10*^fl/fl^. On postnatal day 5 (P5), mice were administered 50 μg of 4-hydroxy-tamoxifen (H7904, Sigma-Aldrich) by intragastric injection to induce genetic inactivation of the endothelial Pdcd10 gene in littermates with *Slco1c1-CreERT2;Pdcd10*^fl/fl^ (*Pdcd10*^BECKO^), and *Pdcd10*^fl/fl^ mice were used as littermate controls. All animal experiments were performed in compliance with animal procedure protocols approved by the University of California, San Diego Institutional Animal Care and Use Committee.

### Brain endothelial isolation

Adult P70 *Pdcd10*^BECKO^ mice and *Pdcd10*^fl/fl^ control littermates were sacrificed, and their brains were isolated and placed into cold solution A (0.5% bovine serum albumin (BSA) in DMEM and 1 μg/μL glucose, 10 mM HEPES, 1x penicillin-streptomycin). Meninges and choroid plexus were removed and minced using scissors in cold solution A. We used two brains of *Pdcd10*^BECKO^ mice and four brains of *Pdcd10*^fl/fl^ mice, which were pooled together, respectively, to collect enough microvasculature. Brain tissue suspension was then centrifuged at 1000g for 5 minutes at 4 °C. The supernatant was removed, and the tissue was digested with a collagenase/dispase solution (1 mg/ml collagenase/dispase (Sigma-Aldrich, #11097113001), 20 units/ml DNase I (Sigma-Aldrich, #DN25), and 0.150 μg/ml tosyl-lysine-chloromethyl ketone (Sigma-Aldrich, #T7254) in DMEM]) at 37 °C for 1 hour with vigorous shaking every 10 minutes. Tissue suspension was triturated using thin-tipped Pasteur pipettes until fully homogenous and centrifuged at 700 g for 5 minutes at 4 °C. The supernatant was removed, and the pellet was resuspended in 20 mL of 25% BSA solution followed by centrifugation at 1000 g for 20 minutes at 4 °C. Capillary fragments were pulled down to the bottom of the tube, remaining BSA and myelin were discarded, and the pellet was resuspended in cold solution A, followed by centrifugation at 700 g for 5 minutes at 4 °C. The supernatant was removed, and capillary fragments were incubated in collagenase/dispase solution at 37 °C for 1 hour. Solution A was added to inactivate enzymatic activity, and the suspension was centrifuged once at 700g for 5 minutes at 4 °C. The cell pellet was resuspended in ACK lysis buffer (Gibco) to lyse red blood cells, and then the cells were centrifuged at 700 g for 5 minutes at 4 °C. The supernatant was removed, and cells were then incubated with anti-CD45-coated beads and passed through a column, following the manufacturer’s protocol (Miltenyi Biotec). Isolated BECs were recovered by negative selection, centrifuged at 700 g for 5 minutes at 4 °C. Pellet was resuspended in frozen media (FBS + 10% DMSO) and stored in liquid nitrogen.

### Nucleus preparation from cryopreserved brain endothelial cells for Chromium single-cell multiome ATAC and gene expression analysis

Brain endothelial cells were thawed according to the 10x Genomics thawing protocol for primary cells (CG000365-Rev B). Briefly, cryovials were removed from liquid nitrogen storage and thawed in a 37°C water bath for 2 min. The thawed cells were transferred to a 50-ml conical tube, and cryovials were rinsed with 1 ml pre-warmed media (DMEM low glucose + 1% FBS). The rinse was added dropwise to the 50-ml conical. Cells were then gradually diluted by five sequential 1:1 volume additions of pre-warmed media, waiting 1 minute between additions and adding at a rate of 1 mL/5 sec. After dilution, cells were centrifuged at 300 g for 5 min. The supernatant was reduced to 1 mL, and the pellet was gently resuspended in this volume. An additional 9 ml of pre-warmed media was added to achieve a total volume of 10 ml and centrifuged at 300 g for 5 min. The supernatant was carefully removed, and the cell pellet was resuspended in 1 mL PBS + 0.04% BSA. The suspension was transferred to a 2-ml microcentrifuge tube, and the 50-ml conical tube was rinsed with 0.5 ml PBS + 0.04% BSA, which was then added to the microcentrifuge tube. After gently inverting it to mix, the cells were centrifuged once more at 300 g for 5 minutes. The supernatant was removed, and the pellet was resuspended in 1 mL PBS + 0.04% BSA for subsequent nuclei isolation.

### Nuclei preparation for Chromium Single-nucleus Multiome ATAC + Gene Expression (10x Genomics)

Brain endothelial cells were permeabilized in lysis buffer (10mM Tris-HCl pH 7.4, 10mM NaCl, 3mM MgCl_2_, 0.01% Tween-20, 0.01% IGEPAL, 0.001% Digitonin, 1% fatty acid-free BSA in PBS, 1mM DTT, 1U/μL Recombinant RNase inhibitor (Promega N2511), 1X Protease Inhibitor (Roche Complete EDTA-free). Nuclei were incubated on ice for 1 minute, then centrifuged (500 rcf, 5min at 4C) in a swinging bucket centrifuge. Supernatant was discarded and 650uL of Wash Buffer (10mM Tris-HCl pH 7.4, 10mM NaCl, 3mM MgCl2, 0.1%.Tween-20, 1% fatty acid-free BSA in PBS, 1mM DTT, 1U/μL Recombinant RNase inhibitor, 1X Protease Inhibitor) was added without disturbing the pellet followed by centrifuging (500 rcf, 5 min at 4°C) in a swinging bucket centrifuge. Supernatant was removed, and the pellet was resuspended in 7uL of 1X Nuclei Buffer (Nuclei Buffer (10x Genomics), 1mM DTT, 1 U/μL Recombinant RNase inhibitor). 1μL was used for counting on a hemocytometer after staining with Trypan Blue (Invitrogen, T10282). 18,000 nuclei were used for tagmentation reaction and 10x Genomics controller loading. Then libraries were generated following the manufacturer’s recommended protocol (https://www.10xgenomics.com/support/single-cell-multiome-atac-plus-gene-expression). 10x multiome ATAC-seq and RNA-seq libraries were paired-end sequenced on a NextSeq 2000, to assess data quality. If data quality was satisfactory, libraries were deeply sequenced on a NovaSeq 6000 to a target depth of ∼50,000 raw reads per cell for each modality.

### RNA isolation

Total RNA from hCMEC/D3 cells was isolated by TRIzol method according to the manufacturer’s instructions (ThermoFisher Scientific). 1ml of TRIzol was used to homogenize the tissue by passing it through a syringe several times. The lysates were transferred to Phase Lock Gel 2 mL tubes, and 200 μL of chloroform (ThermoFisher Scientific) was added to each tube, mixing vigorously for 15 seconds, followed by incubation at room temperature for 3 minutes. Samples were then centrifuged at 12000 g for 10 minutes at 4 °C, and the aqueous phases containing RNA were collected and transferred to 1.5 mL DNAse/RNAse-free microfuge tubes. To precipitate RNA, 500 μL of isopropanol was added, samples were resuspended and incubated for 10 minutes at room temperature, followed by centrifugation at 12000g for 10 minutes at 4 °C. The supernatant was removed, and the pellet was washed with 1 ml of 75% ethanol followed by centrifugation at 7500 g for 5 minutes at 4 °C. RNA was resuspended in water, and the quantity (ND-1000 spectrophotometer; NanoDrop Technologies) and quality (TapeStation; Agilent) of total RNA were analyzed.

### 10x multiome sequence data processing and clustering

Raw sequencing data were processed using cellranger-arc (10x Genomics), generating single-nucleus RNA-seq (snRNA-seq) UMI count matrices for intronic and exonic reads mapping in the sense direction of a gene. We performed unsupervised clustering with RNA UMI counts using the Seurat (V.5)(*78*) standard analysis pipeline. First, cells were filtered for low-quality nuclei by requiring ≥ 500 ATAC fragments and ≥ 200 genes detected per nucleus. Counts were normalized using *NormalizeData* function. We used *FindVariableGenes* function to identify 2,000 variable genes used for principal component analysis (PCA). Putative multiplets were predicted using DoubletFinder(*79*), and 10% of cells were removed from each sample that had the highest doublet score. Batch correction across samples was performed using CCA integration with the *IntegrateLayers* function from Seurat. A *k*-nearest neighbour graph was built using the first 20 PCs, and clusters were identified using Louvain clustering. To visualize the clusters, we applied the Uniform Manifold Approximation and Projection (UMAP) nonlinear dimension reduction technique. Non-endothelial contaminant nuclei were removed by filtering cells with a normalized expression >2 of *Gfap* (Astrocytes), *Col1a1* (fibroblasts), *Acta2* (smooth muscle cells), and *Ptprc* (immune cells). We annotated brain endothelial cell subtypes by reference mapping to the published mouse endothelial cell atlas(*80*) using Seurat. We initially clustered brain endothelial cells at high resolution (resolution = 5) to capture fine substructure. Cluster identities were assigned based on the dominant cell type (≥50%) from reference mapping. Clusters with mixed or ambiguous identity were manually resolved or labeled as ‘unknown’ for exclusion. Final cell type labels were used to rename cluster identities. Unknown and choroid plexus clusters were removed for downstream analysis. We obtained a total of 23,238 endothelial cell nuclei, which were used for downstream analysis (Supplemental Fig. 1).

### Genome assemblies and annotations

*Homo sapiens* (human) assembly: hg38, GRCh38 annotation: hg38 Gencode v33; *M. musculus* (mouse) assembly: mm10, GRCm38 annotation: mm10 Gencode vM22.

### ATAC–seq peak calling and filtering

We used MACS2 for ATAC–seq peak calling on pseudo bulk ATAC–seq fragments using the MACS2 command callpeak with the parameters –shift −75 –ext 150 -q 0.05 – call-summits –nomodel -f BED. We extended the peak summit by 249 bp upstream and 250 bp downstream to achieve 500 bp width for every peak. We iteratively merged the peaks called by MACS2 for every brain endothelial cell subtype and condition, keeping the summit with the highest MACS2 score for overlapped regions. Since the number of peaks called in each BEC cell type is related to the sequence depth, which is highly variable due to differences in cell subtype abundance, we converted MACS2 peak scores (−log10[*q*]) to score per million (*81, 82*). Peaks with a score per million ≥2 were retained for each BEC cell type. We further filtered human and mouse peaks by removing those with ENCODE blacklist regions (https://mitra.stanford.edu/kundaje/akundaje/release/blacklists/) of hg38 and mm10 genomes. This resulted in a master peak set of 118,618 peaks used for the CreateChromatinAssay using Signac(*83*) and for all ATAC downstream analysis.

### Differential gene expression analysis and differential chromatin accessibility analysis

Pseudo-bulk count matrices were generated for each brain endothelial cell (BEC) subtype and genotype using Seurat’s *AggregateExpression* function (RNA and ATAC assays). Differential expression (DE) and differential accessibility (DA) were performed using edgeR (v.4.6.1). For DE, genes with an average log_2_CPM > 1.5 were retained. Subtype-specific genes in *Pdcd10*^fl/fl^ mice were identified by comparing each BEC subtype to all others. To asses *Pdcd10*^BECKO^ effects, each *Pdcd10*^BECKO^ BEC subtype was compared to its matched *Pdcd10*^fl/fl^ control. For DA, peak filtering was applied per comparison: union peaks across all five BEC subtypes for *Pdcd10*^fl/fl^, and BEC subtype-specific peaksets (called in both *Pdcd10*^BECKO^ and *Pdcd10*^fl/fl^ samples) for identifying subtype-specific chromatin accessibility changes in *Pdcd10*^BECKO^ mice.

### Zonation of biological processes

Gene sets representing key biological processes were obtained from Gene Ontology (GO) annotations: angiogenesis (GO:0001525), coagulation (GO:0050817), inflammatory response (GO:0006954), hypoxia (GO:0001666), transporters (GO:0005215), and chromatin remodeling (GO:0006338). Additional gene sets included mouse transcription factors from a previously published list(*84*) and endothelial metabolic genes from the mouse endothelial cell atlas(*80*). Zonated genes were identified based on the DE between BEC subtypes, retaining only those with a linked putative enhancer. Heatmaps were generated using scaled log_2_ (CPM+1) for each gene and its associated putative enhancer.

### Transcription factor motif enrichment

We performed known motif enrichment analysis using Homer (v5.1). For all cCREs, we scanned a region of ± 200 bp around the center of the element. We used a custom background for the analysis: for Pdcd10fl/fl, union peaks across all five BEC subtypes were used; for Pdcd10BECKO DA regions compared to *Pdcd10*^fl/fl^, union peaks across all BEC subtypes and genotypes were used. For hCMEC/D3 cells (human brain endothelial cells), union peaks across all conditions were used.

### KEGG enrichment analysis

We performed KEGG enrichment analysis using EnrichR(*85*), querying the KEGG 2019 Mouse database for mouse data and the KEGG 2022 Human database for mouse and human data. Analyses were conducted using custom background gene sets: for RNA, all genes included in the corresponding DE analysis were used; for genes linked to accessible putative enhancers, all genes predicted to have a putative enhancer were used as the background.

### Inducible PDCD10 KD (TRMPV-siRNA-PDCD10-hCMEC/D3) cell generation Cell culture

Human brain endothelial cell line, hCMEC/D3, at passages 23–33, was routinely cultured in EGM-2 Endothelial Cell Growth Medium-2 BulletKit (Lonza, #CC-3162), hereafter referred to as EGM-2 medium. This medium was supplemented with the following components obtained from the manufacturer at specified concentrations: 0.025% (v/v) rhEGF, 0.025% (v/v) VEGF, 0.025% (v/v) IGF, 0.1% (v/v) rhFGF, 0.1% (v/v) gentamicin, 0.1% (v/v) ascorbic acid, 0.04% (v/v) hydrocortisone, and 2.5% (v/v) FBS(*86, 87*). Tissue culture flasks were precoated with a 1:20 collagen type I solution. The cells were then seeded onto the collagen-coated flasks and maintained at 37°C in an atmosphere of 95% air and 5% CO2 until they reached confluence.

### Human CCM in vitro model

We adopted an inducible RNAi system (to knockdown CCM genes) that allows us to identify retrovirally transduced cells and RNAi induction through the expression of two fluorescent reporters, as previously reported(*88*). This approach uses an inducible tetracycline-responsive element (TRE) promoter that controls the expression of a dsRed fluorescent protein and a microRNA-embedded shRNA directed against *PDCD10* and a second promoter, the phosphoglycerate kinase (PGK) that controls the constitutive expression of the yellow-green fluorescent protein Venus(*88*) that we denominated TRMPV-*PDCD10* (TRE-dsRed-miR30-against-PDCD10-PGK-Venus). We generated stable hCMEC/D3 cell lines using this RNAi system, each expressing one of three different TRMPV-*PDCD10* constructs. We observed that hCMEC/D3 cells transduced with retroviral particles of TRMPV-*PDCD10* after purification by FACS and in the presence of 2 μg/mL doxycycline for 20 days produced cells with 98% of TRE-dsRed-miR30-against-PDCD10-PGK-Venus positive.

### In vitro CCM-like environment and AP-1 inhibition

Cells were plated at a density of 2x10^4^/cm^2^ were plated in collagen-precoated wells in Milicell EZ slide 8-well glass (immunofluorescence) (Millipore Sigma, #PEZGS0816), 6-well plates (RNA-seq and ATAC-seq), or 100-mm plates (ChIP-seq) for 48 hours. CCM environment (500 uM DMOG [Cayman Chemical, #71210] and 10 ng/mL TNF [PeproTech, #315-01A]) and AP-1 inhibitor (40 uM T-5224 [Selleckchem, #S8966]) were prepared in reduced EGM-2 medium (0.012% (v/v) rhEGF, 0.012% (v/v) IGF, 0.05% (v/v) rhFGF, 0.1% (v/v) gentamicin, 0.1% (v/v) ascorbic acid, and 2.5% (v/v) FBS). Cells were exposed to the CCM environment for 36 hours. For AP-1 inhibition, cells were pre-treated for 2 hours with T-5224, followed for 36 hours under CCM environment with T-5224. Cells were then processed for immunohistochemistry or nuclei isolation.

### Bulk ATAC sequencing

ATAC-seq was performed as described(*89*) with few modifications. Cells were resuspended in 250μL of OMNI permeabilization buffer (10mM Tris-HCl pH 7.4, 10mM NaCl, 3mM MgCl_2_, 0.1% NP40, 0.1% Tween20, and 0.01% Digitonin) and incubated on ice for 5 minutes. Nuclei were centrifuged for 5min at 500 rcf at 4°C. Pellets were resuspended in 25uL of tagmenetation buffer (33mM Tris-acetate pH 7.8, 66mM K-acetate, 11mM Mg-acetate, 16% DMF). Nuclei were counted on a hemacytometer, and adjusted to 5,000 nuclei/μL in tagmentation buffer. 10μL (50,000 nuclei) were transferred to a 0.2mL tube and 0.5μL of Tn5 (Tagment DNA enzyme 1, FC-121-1030, Illumina), and incubated at 37 °C for 30min at 500 rpm in thermomixer. DNA was then isolated using MinElute PCR Purification Kit (28004, Qiagen) and eluted in 10μL EB. 10μL of eluted DNA was amplified with 8 cycles of PCR with 25μL NEBNext 2x PCR MasterMix (M0541, NEB), 1μM Nextera i7 and i5 primers, and 11μL water. PCR products were purified using MinElute PCR Purification Kit (28004, Qiagen) and eluted in 20μL EB.

### Chromatin immunoprecipitation sequencing (ChIP-seq)

Around 10 million cells were crosslinked in 1% formaldehyde for 10 min followed by quenching with 0.14 M glycine for 30 min at room temperature. Crosslinked cells were lysed in 500μL of ChIP lysis buffer (1% SDS, 50 mM Tris–HCl pH 8, 20 mM EDTA, c0mplete Protease Inhibitor EDTA-free tablet (Roche; Cat#11836153001). Sonication was performed using the Covaris M220 Focused-ultrasonicator. After sonication, 15μg soluble chromatin was diluted 1:10 in ChIP Dilution Buffer (16.7 mM Tris–HCl, 1.1% Triton X-100, 1.2 mM EDTA, 167 mM NaCl), pre-cleared for 1h at 4°C with 20μL of magnetic Protein A/G beads (Pierce; Cat#88803), and incubated overnight with the following amounts of antibodies-7.96μg anti-JUNB (Cell Signaling Technology; Cat#3753), 5μg anti-H3K27ac (Active Motif; Cat#39034), and 2μg anti-H3K4me1 (Abcam Cat#ab8895). Following pre-clearing, 10μL of sonicated chromatin from each condition were saved as input controls. Magnetic Protein A/G beads (Pierce, Cat#88803) were added for an additional 3h of incubation the following day. Bead-immunocomplexes were washed twice for 5 min with each of the following buffers in order: Wash Buffer A (50 mM HEPES pH 7.9, 0.1% SDS, 1% Triton X-100, 0.1% deoxycholate, 1 mM EDTA, 140 mM NaCl), Wash Buffer B (50 mM HEPES pH 7.9, 0.1% SDS, 1% Triton X-100, 0.1% deoxycholate, 1mM EDTA, 500 mM NaCl), LiCl buffer (20 mM Tris–HCl pH8, 0.5% NP-40, 0.5% deoxycholate, 1 mM EDTA, 250 mM LiCl), and TE (50mM Tris–HCl pH 8, 1 mM EDTA). Elution was performed in 250 μl of elution buffer (50 mM Tris–HCl pH8, 1 mM EDTA, 1% SDS). ChIP and input samples were reverse crosslinked overnight at 65°C. Samples were treated with RNase A (Qiagen; Cat#1007885) for 1h at 37°C and Proteinase K (Invitrogen; Cat#25530015) treated for 2 h at 56°C the following day. DNA was extracted using Zymo DNA Clean & Concentrator-5 (Zymo; Cat#D4014). ChIP-seq libraries were constructed with 20ng DNA using the VAHTS Universal DNA Library Prep Kit for Illumina V4 (Vazyme; Cat#ND610-02) and IDT for Illumina Truseq DNA UD Indexes v2 (Illumina; Cat#20042113). Each ChIP-seq was performed with two biological replicates, where starting chromatin came from separate cultures for each replicate. Libraries were sequenced on the illumina NextSeq 2000 P2-200 kit with 101+8+8+101 cycles.

### Identification of orthologous sequence elements across mouse and human

We identified orthologous sequences for each 500bp human or mouse *cis*-regulatory element in the corresponding species using liftOver(*90*). For each element, we first performed liftOver to the other species’ genome (mm10 or hg38) with a requirement of 50% retained sequence identity (minMatch = 0.5) and retained only orthologous elements that are 1 kb or less to the lifted-over genome. We next performed liftOver from the identified orthologous sequence back to the original species. We retained all sequences that overlapped the original location when mapped back to the starting genome.

### Immunohistochemistry

Brains from Adult P70 *tdtTomato*;*Pdcd10*^BECKO^ and littermate control *tdtTomato;Pdcd10*^fl/fl^ mice were isolated and fixed in 4% paraformaldehyde (PFA) in PBS at 4 °C overnight. The tissue was cryoprotected in a 30% sucrose solution in PBS, then embedded and frozen in O.C.T. compound (Fischer Scientific). Brains were cut into 18-μm sagittal sections using a cryostat and placed onto Superfrost Plus slides (VWR International, #1255015). Sections were incubated in a blocking-permeabilization solution (0.5% Triton X-100, 5% donkey serum, 0.5% BSA, in PBS) for 2 hours and incubated in rabbit monoclonal antibodies against JUNB (1:200, #3753S; Cell Signaling), Iba1 (1:100; 019-19741; FUJIFILM Wako) and GFAP (1:300; GA524; Agilent Dako) in PBS at room temperature overnight. Tissue sections were washed four times in PBS and incubated with Alexa Fluor 488 anti-rabbit secondary antibodies (1:300, #711-546-152; Jackson Laboratory) in PBS for 2h at RT. Cell nuclei were stained and mounted using DAPI Fluoromount-G mounting medium (SouthernBiotech, #0100). Human CCM tissue was similarly processed for immunofluorescence analysis, and sections were incubated in rabbit monoclonal antibodies against JUNB (1:200, #3753S; Cell Signaling), goat polyclonal antibody against CD31 (1:100, #AF3628; R&D Systems) in PBS at room temperature. Tissue sections were washed four times in PBS and incubated, with Alexa Fluor 488 anti-rabbit secondary antibodies (1:300, #711-546-152; Jackson Laboratory) and Alexa Fluor 594 anti-goat secondary antibodies (1:300, #705-585-003; Jackson Laboratory) in PBS for 2h at RT. Cell nuclei were stained and mounted using DAPI Fluoromount-G mounting medium. For both mouse and human samples, the slides were viewed with a high-resolution slide scanner (Olympus VS200 Slide Scanner), and the images were captured with VS200 ASW V3.3 software (Olympus). The image processing was performed using ImageJ Ver. 1.53f on high-resolution images.

### Human gene-cCRE correlation analysis

We identified putative enhancer-target gene regulatory pairs in human endothelial cells by finding pairs with correlated activity between conditions. We calculated Pearson correlation coefficients using average CPM of ATAC-seq or RNA-seq across 3 replicates for 8 conditions of treatments (PDCD10 WT+T-5224; PDCD10 KD+T-5224; PDCD10 WT+vehicle; PDCD10 KD+vehicle; PDCD10 WT+CCM; PDCD10 KD+CCM; PDCD10 WT+T-5224+CCM; PDCD10 KD+T-5224+CCM), for every gene and cCRE pair within 500kb between the gene TSS and cCRE coordinates. Pearson correlation p-values were converted to FDR adjusted p-values. Pairs with positive coefficient and an FDR < 0.1 were considered putative enhancer-target gene pairs.

## Results

### Epigenetics significantly influences the transcriptional programs of arteriovenous endothelial cells in the mouse brain vasculature

Single-cell and single-nuclear transcriptomic approaches (scRNA-seq and snRNA-seq)(*91, 92*) have advanced our understanding of arteriovenous transcriptional programs in humans and mice(*93-97*). Currently, there is limited knowledge of the gene regulatory mechanisms driving endothelial cell identity along the brain arteriovenous axis. Additionally, the identity and role of transcription factors in maintaining the arteriovenous endothelial cells is not yet fully understood. We conducted multi-omic analyses to simultaneously profile the brain endothelial transcriptome and chromatin accessibility using 10x Genomics single-nuclei high-throughput snRNA-seq and single-nucleus transposase-accessible chromatin sequencing (snATAC-seq). Our goal was to determine whether arteriovenous endothelial cells exhibit unique chromatin accessibility patterns associated with cell type-specific gene expression (Fig. 1A). For this study, we isolated and highly enriched brain endothelial cells (BEC)(*45, 47*) from adult P70 (Postnatal day 70) male and female *Pdcd10^fl/fl^* healthy mice(Fig. 1A). After removing doublets using DoubletFinder and applying quality filtering, we excluded clusters of contaminating immune cells (marked by *Ptprc*, the gene encoding CD45), smooth muscle cells (*Acta2*), astrocytes (*Gfap*), fibroblasts (*Col1a1*), perycites (*Myh11*, *Pdgfrb*, *Vtn*), and red cells (*Hbb-bs*) (Supplemental Fig. 1). The data were then integrated using canonical correlation analysis (CCA) included in the Seurat package(*78*) to correct for batch effects and performed unsupervised clustering based on gene expression. After quality control and exclusion of non-endothelial cells, we annotated subtypes of BEC based on reference mapping genes from published transcriptome datasets(*80, 93-97*) (Fig. 1B,1C). BEC were categorized into five subtypes: artery (*Fbln5*, *Gkn3, n =* 368), capillary arterial (*Tgfb2, n =* 796), capillary (*Mfsd2a, Rgcc, n =* 5,401), capillary venous (*Tfrc*, *Slc16a1, n =* 6,110), and large vein (*Vwf, Vcam1, n = 572*) (Fig. 1B-1D), and found proportions of each subtype to be highly consistent across biological replicates (Fig. 1C). As previously reported, we observed that the BEC subtype has distinctive transcriptional gene programs that contribute to vascular heterogeneity in the mouse cerebral vasculature(*93, 95, 98*) (Fig. 1D, 1E).

**Figure 1.**
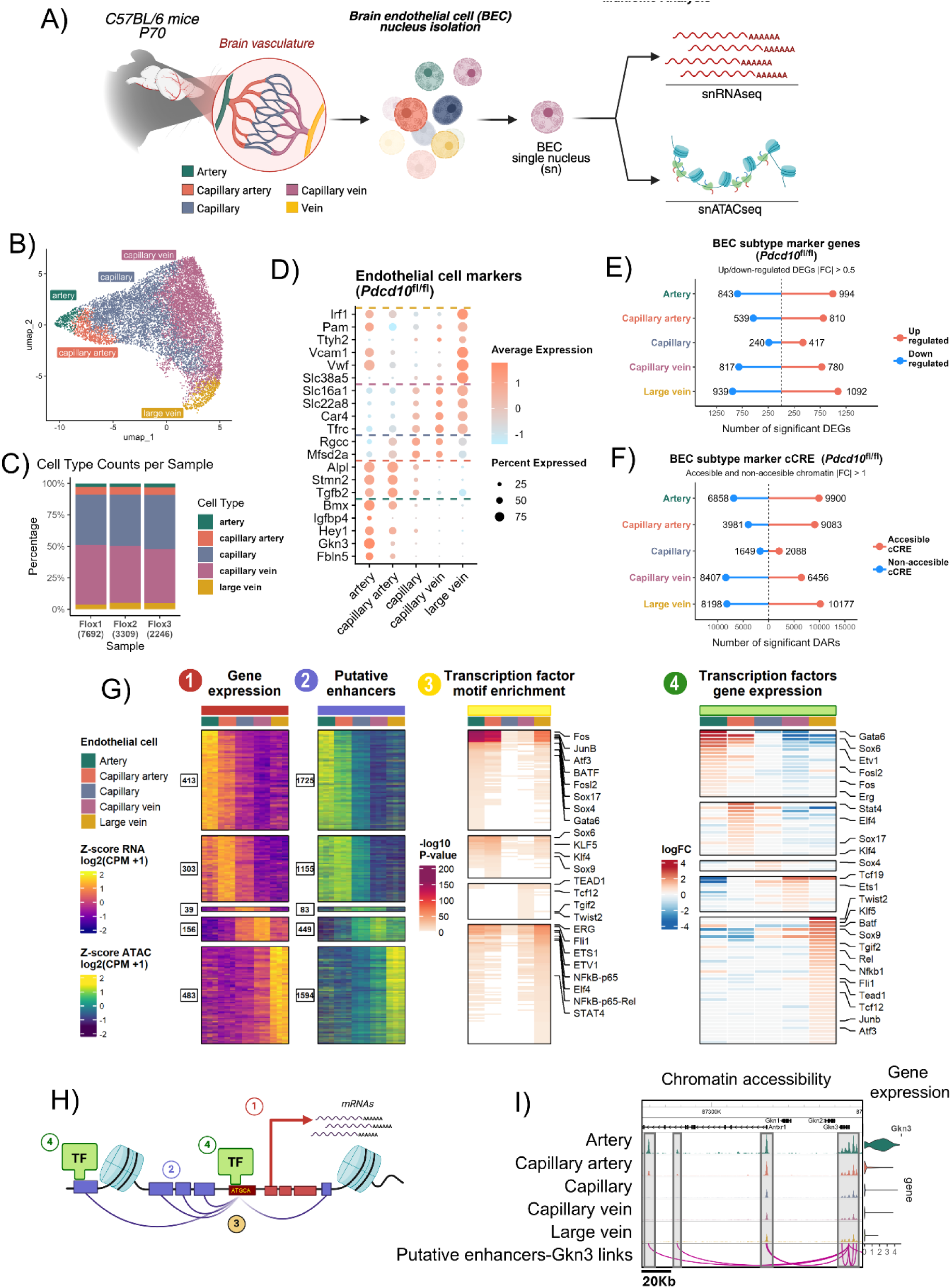
Epigenetics influences the transcriptional programs of endothelial cells in the brain arteriovenous axis. **A)** Schematic illustration of the experimental design. Brain vasculature was isolated from pooled samples of two animals per replicate for *Pdcd10*^BECKO^ (n = 3) and *Pdcd10*^fl/fl^ (n = 3) mice. Nuclei were isolated and processed using 10x Genomics Multiome for simultaneous snRNA-seq and snATAC-seq. **B)** Uniform manifold approximation and projection (UMAP) clustering with annotation of distinct brain endothelial cell (BEC) subtypes in *Pdcd10*^fl/fl^ mice. **C)** Distribution of BEC subtype (y-axis) across *Pdcd10*^fl/fl^ mouse replicates. Bar labels indicate the total number of nuclei per replicate. Colors correspond to the five BEC subtypes. **D)** Dot plot shows the average expression levels of BEC subtype markers and percentage of expressing cells. Dashed lines separate the five different BEC subtypes. **E)** Number of differentially expressed genes (DEGs) from each BEC subtype compared with the rest. Red lines indicate upregulated genes; blue lines indicate downregulated genes. Significance is defined as |log_2_ fold change (FC)| > 0.5 and FDR < 0.05. **F)** Number of differentially accessible regions (DAR) from each BEC subtype compared with the rest. Red lines indicate regions with gained chromatin accessibility (CA); blue lines indicate lost CA. Significance is defined as |log_2_ FC| > 1 and FDR < 0.05. **G)** Integrated heatmaps summarizing transcriptional and epigenetic relationships across BEC subtypes. *1)* Heatmap showing gene expression (Z-score of RNA log_2_ [CPM + 1]) for BEC subtype-specific marker genes. Numbers on the left indicate the number of identified marker genes per subtype. *2)* Heatmap showing chromatin accessibility of putative enhancer regions (Z-score of ATAC accessibility log_2_ [CPM + 1]) linked to the same marker genes. Numbers on the left indicate the number of identified marker putative enhancers per BEC subtype. Each row in (1) and (2) represents a gene-enhancer pair. *3)* HOMER transcription factor (TF) motif enrichment analysis showing - log_10_ *p*-values for putative enhancers identified in (1). Representative enriched TF motifs are labeled. *4)* Heatmap showing predicted TF expression as log_2_ FC across BEC subtypes, corresponding to the TFs identified in (3). **H)** Schematic summarizing the integrative analysis in (G), including: 1) gene expression, 2) enhancer accessibility, 3) TF motif enrichment, and 4) TF gene expression. **I)** WashU Epigenome Browser snapshot of aggregate chromatin accessibility around the *Gkn3* locus across BEC subtypes. **Bottom:** arcs represent predicted links between the *Gkn3* transcription start site (TSS) and putative enhancer peaks, with a 20-kilobase scale bar indicating enhancer–gene distances. **Right:** violin plots show *Gkn3* expression across BEC subtypes.

We next analyze the snATAC-seq modality to create an atlas of chromatin accessibility for each BEC subtype (Supplemental Fig. 1). Chromatin-accessible genomic regions were identified through peak calling using MACS2(*99*) in each subtype. Overlapping peaks across subtypes were merged to create a union, non-redundant atlas of 125,301 candidate cis-regulatory elements (cCREs) across BEC subtypes. This revealed BEC-specific cCREs at non-coding promoter distal and promoter proximal regions of endothelial-specific markers (Supplemental Fig. 1). An average of 82,042 BEC-specific open chromatin regions in *Pdcd10^fl/fl^* mice were identified. We generated pseudo-bulk datasets by aggregating counts for each subtype of endothelial cells, along with their corresponding chromatin accessibility and gene expression for downstream analysis. Differentially accessible chromatin regions (DARs) were identified, which were either “accessible cCRE” or “non-accessible cCRE” in each BEC subtype (Fig. 1F). We found that arteries, capillary arteries, and large veins have a higher number of accessible cCREs (∼9,000) than capillaries and capillary veins (∼5,000) (Fig. 1F). Notably, arteries, capillary veins, and large veins display significantly larger regions of non-accessible cCRE (∼8,000) compared to capillary arteries and capillaries (∼2,000) in adult mice (Fig. 1F). These findings reveal BEC subtype-specific gene and cCRE markers.

We next identified putative enhancers and their target genes in BEC through the LinkPeaks function from Signac(*83, 100*), which correlates normalized chromatin accessibility signals with RNA expression for each pair of distal cCRE/putative enhancers and gene promoter within 500 kilobases(*100-103*). This analysis yielded 34,453 positively correlated putative enhancer–target gene pairs with an empirically defined significance threshold of p-value <0.05 (Fig. 1G-1I), with a median distance between the potential enhancers and the target promoters of 125 kb. The putative enhancer-target gene pairs revealed BEC subtype-specific gene regulatory programs in arteries (413 marker genes and 1,725 putative enhancers), capillary arteries (303 marker genes, 1,155 putative enhancers), capillary (39 marker genes, 83 putative enhancers), capillary vein (156 marker genes, 449 putative enhancers), and large veins (483 marker genes, 1594 putative enhancers) (Fig. 1G). To predict the transcription factors (TFs) driving BEC subtype-specific gene regulatory programs, TF motif enrichment analysis was performed using HOMER software(*104*). Unique TF motifs enriched in the BEC subtype-specific putative enhancers were identified, many of which had matching TF expression patterns across the subtypes (Fig. 1G). For example, arteries showed the strongest enrichment for Activator Protein-1 (AP-1) family members (e.g., *Fos, Fosl2, Etv1*), whereas large veins had the strongest enrichment for the Nuclear factor kappa B (NfκB) family (e.g., Nfkb1, Rel). The Erythroblast Transformation Specific (ETS) family was also enriched along the arteriovenous axis with different members enriched in the different BEC subtypes (e.g., Erg, Etv1, Ets1, Fli1) (Fig. 1G). The chromatin accessibility profiling of each arteriovenous BEC subtype revealed their epigenomes to be highly diverse, suggesting a major influence of epigenetic regulation on subtype-specific transcriptional programs. Analysis of these subtype-specific transcriptional programs identified putative TFs critical for establishing BEC subtype identities and functions.

### Epigenetics significantly influences the transcriptional programs linked to the zonation of biological processes in brain endothelial cells

In the brain arteriovenous axis, the expression of arterial and venous marker genes peaked at opposite ends of this range. The gradual changes in the expression of these markers in smaller endothelial cells suggest a zonation in the expression of TFs, which confers specialized functions and specific morphological characteristics(*97, 98, 105*). We observed zonation-dependent gene expression in the BEC subclusters of transcription factors (113 genes) and transmembrane transporters (95 genes) as previously reported(*106*), and additional genes for relevant biological processes such as genes involved in endothelial metabolism (34 genes), inflammatory response (123 genes), angiogenesis (103 genes), hypoxia signaling (50 genes), coagulation processes (38 genes), autophagy (50 genes), and chromatin remodeling (95 genes) (Fig. 2A, Supplemental Fig. 2). Additionally, single-nucleus chromatin accessibility revealed putative enhancers with accessibility that correlated with transcriptomic levels of BEC zonation genes in relevant biological processes for each endothelial subtype (Fig. 2A, Supplemental Fig. 2). Furthermore, TF motif enrichment analysis showed significant enrichment of binding motifs for the AP-1, ETS, KLF, SOX, Mef2, and TEAD families in putative enhancers/cCRE that become more accessible in BECs during the zonation of key endothelial biological processes (Fig. 2B), suggesting that these TFs may represent core regulators of brain endothelial zonation. Furthermore, we identified process-specific enrichment of TF binding motifs (Fig. 2C), indicating the presence of complementary transcriptional regulators that may drive the unique zonation profiles linked to particular biological processes specific to BEC subtypes. This suggests that the zonation of the brain arteriovenous axis in the mouse brain is influenced by epigenetic regulation and particular transcription factors in the brain endothelium.

**Figure 2.**
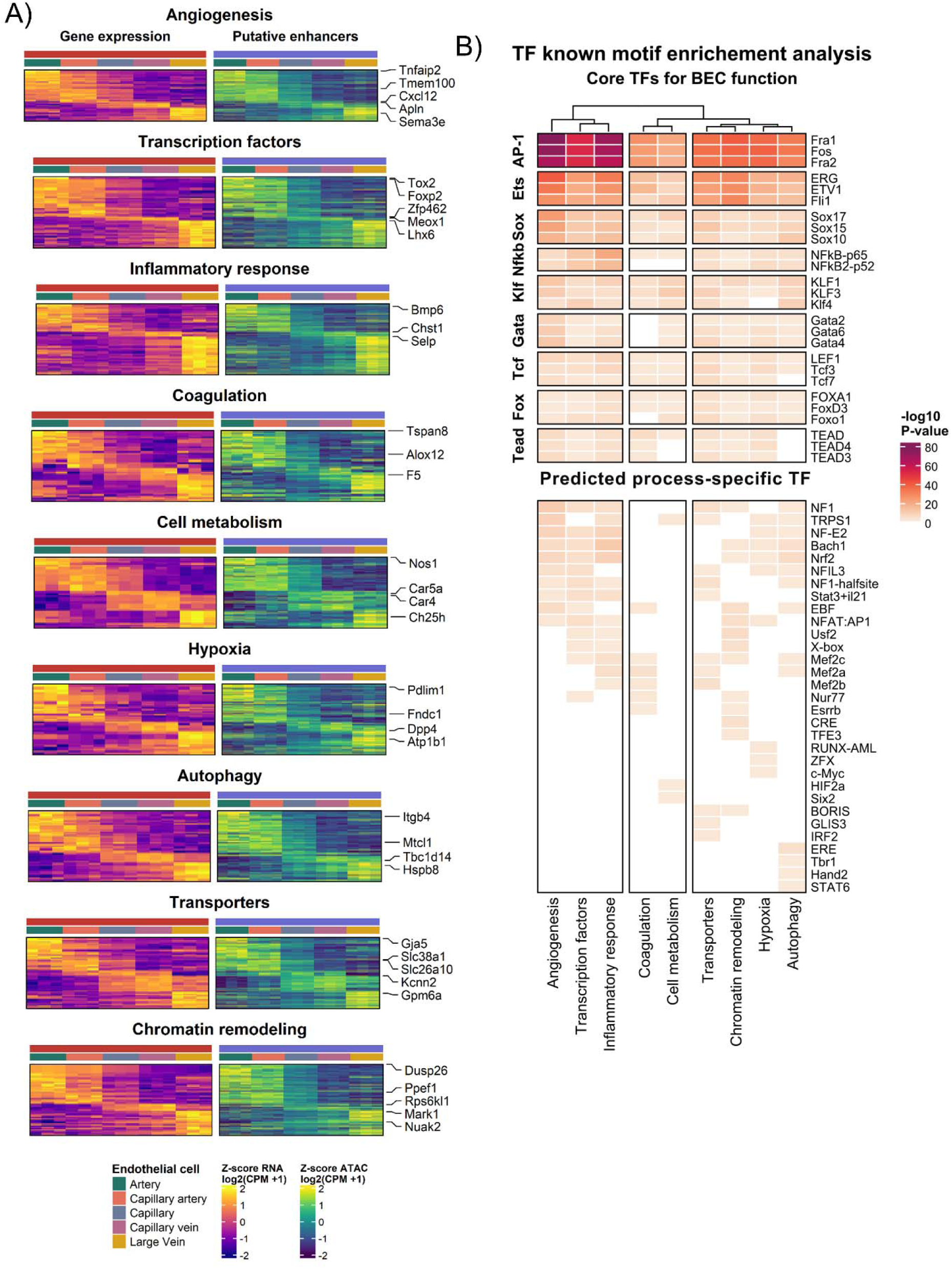
Epigenetics influences the zonation of biological processes in brain endothelial cells. **A)** Heatmap showing gene expression and corresponding putative enhancer accessibility for zonated genes across processes. Left: Z-score of RNA expression (log_2_ [CPM + 1]); right: Z-scores of ATAC accessibility (log_2_ [CPM + 1]). Each row represents a gene-enhancer pair. Representative genes are labeled. **B)** HOMER motif enrichment analysis showing -log_10_ *p* values for putative enhancers identified in (A). Top: motifs enriched across all processes, representing core transcription TF for BEC function. Bottom: predicted process-specific complementary TFs.

### Epigenetics and gene expression reveal a persistent activated state of endothelial cells in CCM disease

A growing body of evidence indicates that the activation of BEC due to neuroinflammation significantly contributes to the chronic nature and worsening of CCM disease and its associated morbidity(*13, 46, 49, 107-109*). However, the underlying inflammatory mechanisms have yet to be entirely understood. To understand how neuroinflammation affects BECs during CCM disease, we investigated brain endothelial gene expression patterns and chromatin accessibility in adult *Pdcd10^BECKO^*mice, a chronic animal model of CCM disease(*45, 47*), and compared findings with *Pdcd10^fl/fl^* littermate controls (Fig. 1-3, Supplemental 1-3). Consistent with our previous study of RNA-seq profiling of fresh BECs isolated from *Pdcd10^BECKO^*brains and *Pdcd10^fl/fl^* control brains, extensive changes were observed in gene expression in BECs during CCM disease compared to controls, associated with inflammation, thrombosis, and endothelial dysfunction(*45, 47*). Notably, among the BEC subtypes, the capillary vein cluster exhibited the most significant and extensive changes in gene expression during CCM disease (Fig. 3A, Supplemental Fig. 3). We found up-regulation of 2,137 genes and down-regulation of 1,625 genes in *Pdcd10^BECKO^* mice (Fig. 3A). In contrast, the artery cluster had relatively milder changes in gene expression with up-regulation of 385 genes and down-regulation of 152 genes (Fig. 3A). The capillary and large vein subclusters also showed an extensive gene expression changes during CCM (Fig. 3A). Importantly, 549 genes were specifically upregulated in the large vein cluster, 563 genes showed specific upregulation in the capillary-to-large vein axis, and 160 genes were upregulated along the arteriovenous axis in *Pdcd10^BECKO^* mice (Supplemental Fig. 3). These results are consistent with previous studies in which CCM disease primarily affects veins(*18*). We then analyzed DARs in *Pdcd10^BECKO^*to determine accessible or non-accessible cCREs in the BEC subtypes when compared to BEC subtypes in littermate control *Pdcd10^fl/fl^* (Fig. 1, Fig. 2). We observed that most DARs correspond to increased chromatin accessibility (∼70%) in BEC subtypes from *Pdcd10^BECKO^* mice (Fig. 3B). Similar to gene expression, the large vein cluster exhibited the highest number of DARs, with 18,263 regions gaining accessibility and 7,775 regions losing chromatin accessibility. Arteries showed less extensive changes, with only 1,235 regions gaining chromatin accessibility and 415 regions losing it. Notably, the capillary vein cluster had the most DARs among the capillary subtypes, with 19,305 regions gaining accessibility and 6,697 regions losing accessibility. In addition, 5,170 regions gained chromatin accessibility specifically in the large vein cluster, 5,144 gained chromatin accessibility specifically in the capillary-to-large vein axis, and 5,073 specifically gained accessibility in capillaries and large vein in *Pdcd10^BECKO^*mice (Supplemental Fig. 3). These findings demonstrate large-scale epigenomic reprogramming in BEC subtypes during CCM disease.

**Figure 3.**
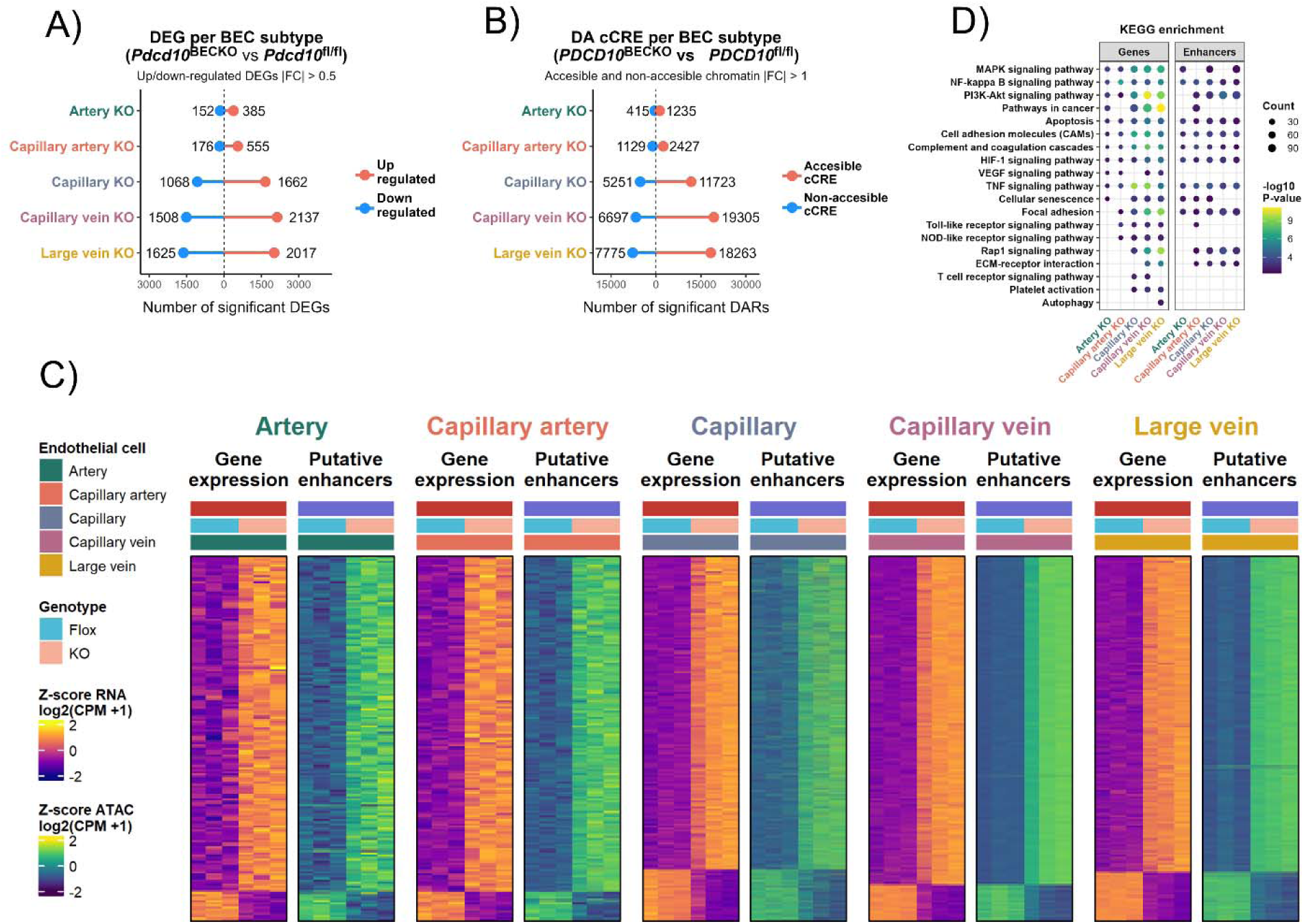
Reprogramming of the brain endothelial cell epigenome is linked to inflammation, thrombosis, and endothelial dysfunction in *Pdcd10^BECKO^* mice. **A)** Number of DEGs between *Pdcd10*^BECKO^ and *Pdcd10*^fl/fl^ in each BEC subtype. Red lines indicate upregulated genes; blue lines indicate downregulated genes. Significance is defined as |log_2_ FC| > 0.5 and FDR < 0.05. **B)** Number of DARs between *Pdcd10*^BECKO^ and *Pdcd10*^fl/fl^ in each BEC subtype. Red lines indicate regions with gained CA; blue lines indicate lost CA. Significance is defined as |log_2_ FC| > 1 and FDR < 0.05. **C)** Integrated heatmaps summarizing transcriptional and epigenetic changes between *Pdcd10*^BECKO^ and *Pdcd10*^fl/fl^ across BEC subtypes. For each subtype: left panels show gene expression of up- and downregulated genes; right panels show chromatin accessibility of linked putative enhancers. Gene expression and chromatin accessibility are shown as row-wise Z-scores of log_₂_ (CPM + 1). Z-scores were calculated independently for each gene (RNA) and each enhancer (ATAC) within each BEC subtype, across genotypes. Each row represents a matched gene–enhancer pair. **D)** KEGG pathway enrichment analysis of up-regulated genes and more accessible putative enhancers in *Pdcd10*^BECKO^ across BEC subtypes. Color scale represents -log10 *p*-values; circle size indicates the number of genes per term.

Moreover, we evaluated the influence of the BEC cCRE/putative enhancers on gene expression by assessing the change in expression of their predicted target genes between *Pdcd10^BECKO^*and litter controls *Pdcd10^fl/fl^* mice (Fig. 3C). This analysis identifies putative enhancers that are critical for gene expression in various brain endothelial subtypes during CCM disease. Putative enhancers were mostly located in promoter-distal regions, with 57.4% located in introns and 32% in distal intergenic regions (Supplemental Fig. 3). We used all upregulated genes as input, and genes linked to increased accessible putative enhancers identified within each BEC subtype, then conducted a KEGG(*110*) enrichment analysis with EnrichR(*85, 111, 112*)by integrating multi-omic data from snRNA-seq and snATAC-seq to identify common regulatory pathways influenced by changes in chromatin accessibility and gene expression. The analysis revealed significant enrichment for terms related to inflammation, hypoxia signaling, PI3K-Akt pathway, cellular senescence, Rap1 signaling, among others (Fig. 3D). These findings indicate that epigenomic reprogramming in CCM disease promotes signaling pathways associated with a persistently activated endothelial cell state. Notably, vascular inflammation and thrombosis are linked to CCM disease-driven morbidity and chronicity of the disease(*47*). We observed that BEC from *Pdcd10^BECKO^* mice showed an increase in chromatin accessibility at the promoter and promoter distal regions of CCM-associated genes involved with leukocyte recruitment: *Vcam1*, *Icam1*, *Cd40*, *Sele* (E-selectin), and *Selp* (P-selectin), and genes associated with inflammation and thrombosis: *Cx3cl1*, *PAI-1(Serpine1), EPCR (Procr), vWF, Factor V*, *Cd74, Tgfb1, Il6*, in an endothelial cell type-specific manner (Fig 4A-4C). Conversely, a cell type-specific decrease in chromatin accessibility was noted upstream of the *tPA* promoter at a putative enhancer among others (Fig. 4A-4C). These changes in chromatin accessibility are directly correlated with transcriptomic levels in the same endothelial cell subtype (Fig. 4B, 4C). Recent studies have demonstrated the significant role of these genes in chronic CCM disease. Specifically, they increase leukocyte recruitment and activation (such as CX3CL1, VCAM1) and modulate fibrinolysis (such as PAI-1, also known as Serpine1, and tPA, also known as Plat)(*45, 47*). They have also been implicated as direct platelet activators and prothrombotic drivers in CCM (such as vWF)(*113*) and neuroinflammation (such as CD74, H2-Ab1) (*45, 47*). These findings suggest that changes in epigenetics of BEC during CCM disease affect gene expression patterns, driving a persistent activated state in endothelial cells linked to leukocyte recruitment and activation, thrombosis, and endothelial dysfunction in CCM pathology (Fig. 4D).

**Figure 4.**
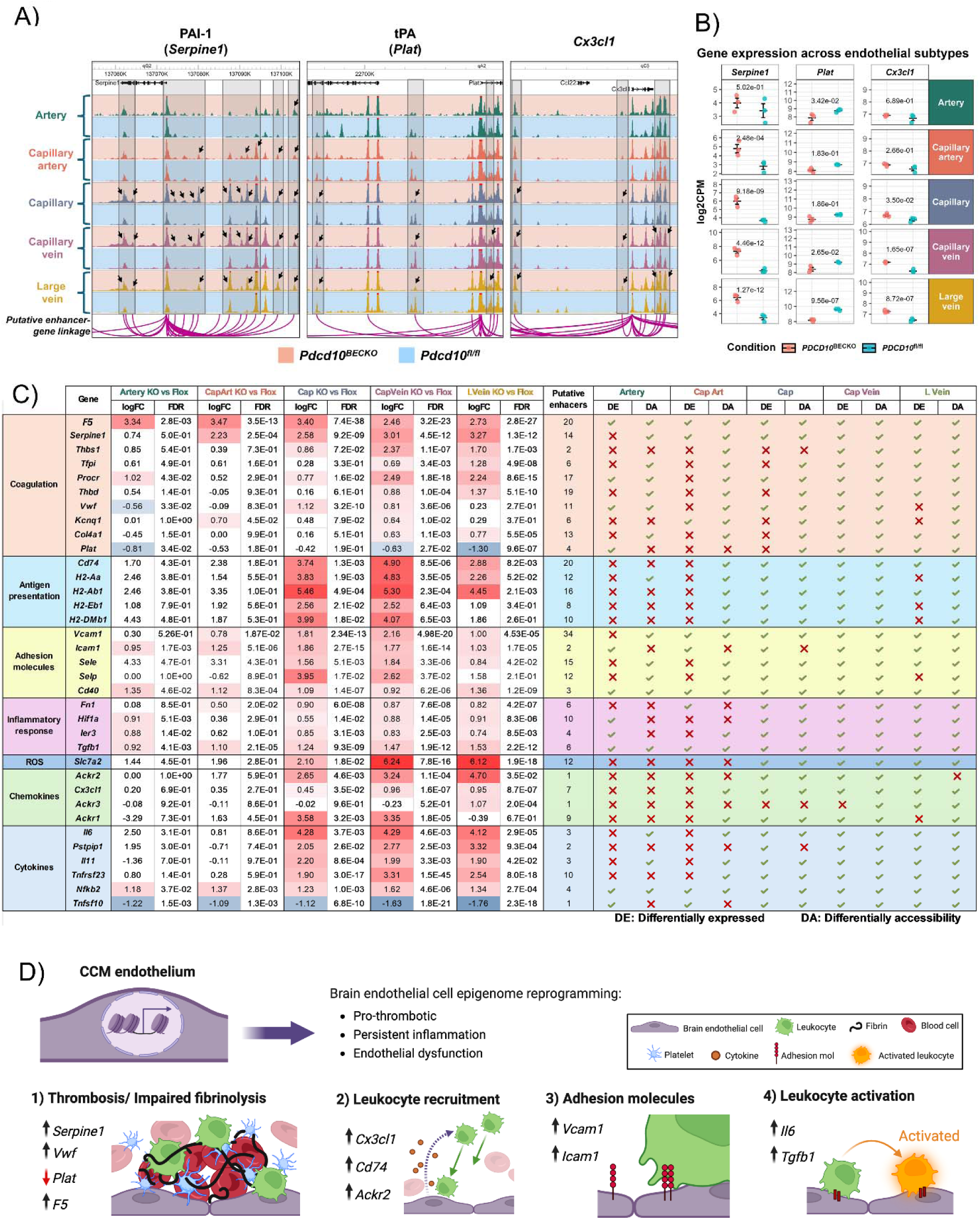
Persistent activated brain endothelial cell state is linked to inflammation, thrombosis, and endothelial dysfunction in *Pdcd10^BECKO^*mice. **A)** WashU Epigenome Browser snapshots showing aggregate chromatin accessibility around the *Serpine1* **(left)**, *Plat* **(middle)**, and *Cxc3cl1* **(right)** loci across BEC subtypes. Arcs represent predicted links between each gene’s transcription start site (TSS) and its associated putative enhancer. Pink background indicates *Pdcd10*^BECKO^; blue background indicates *Pdcd10^f^*^l/fl^. Regions with differential enhancer accessibility are highlighted in gray and indicated by arrows. **B)** Dot plots with error bars showing gene expression (log_2_ [CPM + 1]) of *Serpine1* **(left)**, *Plat* **(middle)**, and *Cxc3cl1* **(right)** across BEC subtype in *Pdcd10*^BECKO^ (pink dots) and *Pdcd10^f^*^l/fl^ (blue dots). Error bars represent mean ± SEM. FDR values were calculated using edgeR’s *glmLRT* function. Panel colors indicate endothelial subtype. **C)** Summary of gene expression, putative enhancers, and chromatin accessibility of genes associated with vascular inflammation and thrombosis. Log_2_ FC and FDR values are shown for each gene across BEC subtype. The number of putative enhancers linked to each gene is indicated. Differential expression (DE) and differential accessibility (DA) are marked with a check mark if significant for each BEC subtype. Capillary arterial (CapArt); capillary (Cap); capillary vein (CapVein); and large vein (LVein). **D)** Schematic illustration summarizing epigenetic changes at inflammatory loci, which lead to a persistent activated state of brain endothelial cells in CCM disease, impacting inflammation, thrombosis, and endothelial dysfunction.

### The AP-1 transcription factor JUNB is an activator of transcriptional programs associated with a persistently activated endothelial cell state in CCM disease

Although it is known that inflammation contributes to the exacerbation of CCM disease(*45, 48, 49*), the specific role of inflammation in persistent activation of endothelial cells, and propensity for thrombosis is not fully understood. We hypothesize that a persistent activated endothelial cell state in CCM disease relies on inflammation-induced chromatin and transcriptomic changes, driven by the activation of specific transcription factors. We began our analysis by examining transcription factor motifs enriched in DARs (Fig. 3D, 3C) in BEC isolated from *Pdcd10^BECKO^* mice (Fig. 5A). Our findings revealed a strong enrichment for binding motifs of the AP-1 transcription factor family, which includes JUNB, BATF, ATF3, FOS, FRA1, FOSL1 (Fig. 5A). Additionally, we observed enrichment for the ETS transcription factor family, comprising ERG, ETV1, ETS1, and ETV4, in the elements that exhibit increased accessibility in BEC from *Pdcd10^BECKO^* mice (Fig. 4A). We further examined TFs that may contribute to the reprogramming of BEC towards a persistently activated endothelial cell state in *Pdcd10^BECKO^* brains. We noticed that *Junb* expression was significantly increased in *PDCD10*^BECKO^ brain endothelium compared to littermate controls in most vascular beds (Fig. 5B). *Junb* has been shown to play an essential role in inflammatory processes, particularly in immune cells(*114, 115*). Moreover, we identified putative JUNB target genes as those linked to cCRE/putative enhancers containing JUNB motifs in BEC from *Pdcd10^BECKO^*mice. JUNB target genes showed significantly increased expression across all brain endothelial subtypes in *PDCD10*^BECKO^ mice (Fig. 5B). Even more strikingly, the chromatin accessibility of their associated cCRE/putative enhancers was also elevated in BEC from *PDCD10*^BECKO^, consistent with enhanced transcriptional activation (Fig. 5B). We also noticed a substantial increase in brain endothelial JUNB expression in areas with chronic lesions associated with thrombosis in both mouse (Fig. 5C) and human (Fig. 5D) tissues. These results suggest that during chronic CCM disease, there is an increased activity of putative enhancers with JUNB motifs, leading to the upregulation of their target genes in CCM endothelial cells. We repeated this analysis for NF-kB1 and found a similar trend, with elevated levels of *Nfkb1* mRNA at different brain endothelial subtypes, upregulation of putative NF-kB1 target genes, and increased accessibility of putative enhancers with NF-kB1 motifs. However, the number of genes and enhancers related to NF-kB1 was ∼60% and ∼80% less, respectively, than those for JUNB (Supplemental Fig. 4). During the assessment of the ETS family member Erg, we discovered that Erg target genes exhibited a significant increase in expression of BEC in *Pdcd10^BECKO^* mice, which correlated with more accessible enhancers (Supplemental Fig. 4). However, we notice that Erg differential expression was restricted to be upregulated in large veins. We also observed differences in *Klf4* expression levels among capillary veins and large veins, which is a known marker for the formation of CCMs(*17, 19, 116-118*). Although we identified upregulation of KLF4 target genes and more accessible associated putative enhancers, the number of enhancers was significantly fewer than those identified for JUNB. Therefore, these findings highlight the potential role of the endothelial AP-1 transcription factor JUNB in the epigenomic reprogramming of BEC, leading to a persistently activated state in CCM disease.

**Figure 5.**
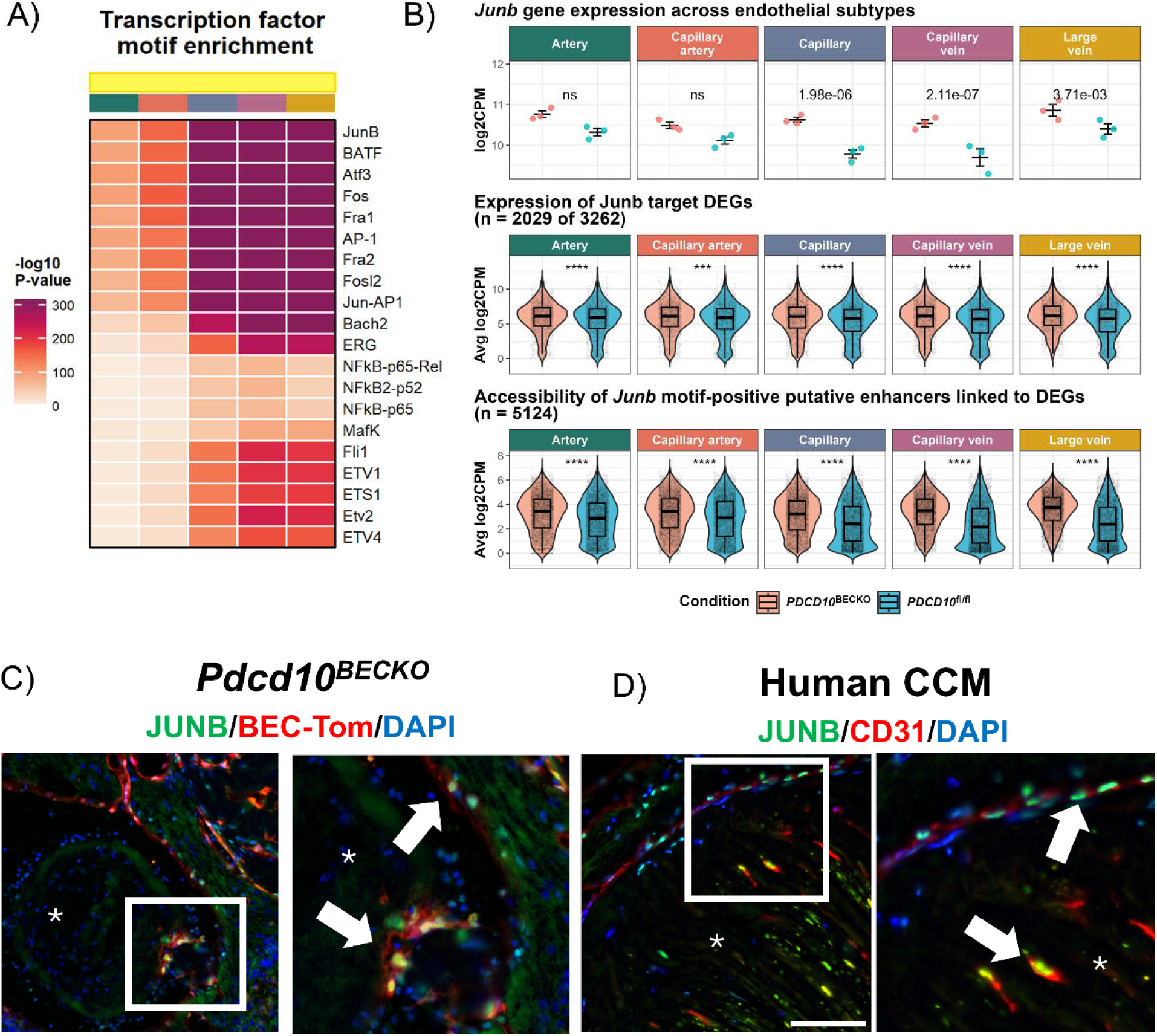
JUNB is an activator of brain endothelial transcriptional programs in CCM disease. **A)** HOMER motif enrichment analysis showing -log_10_ *p*-values for putative enhancers with increased accessibility in *Pdcd10*^BECKO^. **B)** *Junb* expression and regulatory activity across endothelial subtypes. **Top:** *Junb* expression in *Pdcd10*^BECKO^ (pink dots) and *Pdcd10*^fl/fl^ (blue dots) across endothelial subtypes. Each point represents a biological replicate. FDR values were calculated using edgeR’s *glmLRT* function; ns, not significant. **Middle:** Expression of JUNB-regulated differentially expressed genes (DEGs) (n = 2,029 out of 3262) across BEC subtypes. Each point represents one DEG. **Bottom:** Chromatin accessibility of JUNB motif-containing putative enhancers linked to DEGs (n = 5124), per BEC subtype. Each point represents an individual putative enhancer. For panels **(Middle)** and **(Bottom)**, *P*-values from Wilcoxon rank-sum tests. For the **top** panel, error bars represent mean ± SEM. Subtype-specific colors indicate the endothelial subtype. **C)** Immunofluorescence staining in Pdcd10^BECKO^-tdtTomato mice showing colocalization of JUNB (green) with CCM endothelial cells (tdTomato, red), both delimiting and within the CCM lesion. Arrows indicate representative JUNB-positive endothelial cells. Asterisk marks lumen of CCM lesion. Nuclei are stained with DAPI (blue). **D)** Immunofluorescence staining of human CCM lesion showing JUNB-positive (green) CCM endothelial cells stained with CD31 (red). JUNB expression is observed both in the surrounding area and within the CCM lesion. Arrows point to representative JUNB-positive BECs. Nuclei are stained with DAPI (blue).

### The AP-1 transcription factor JUNB contributes to a persistent activated brain endothelial cell state

To study the molecular mechanisms that affect the genomic accessibility that enforced a persistent activated endothelial cell state during chronic CCM disease, we adopted an inducible RNAi system that allows targeting *PDCD10* (CCM3 gene) in a human brain endothelial cell line, hCMEC/D3, in a time-controlled manner. This approach uses an inducible tetracycline-responsive element (TRE) promoter that controls the expression of a dsRed fluorescent protein and a microRNA-embedded shRNA directed against *PDCD10* and a second promoter, the phosphoglycerate kinase (PGK) that controls the constitutive expression of the yellow-green fluorescent protein Venus(*88*) that we denominated TRMPV-*PDCD10* (TRE-dsRed-miR30-against-PDCD10-PGK-Venus). We generated stable hCMEC/D3 cell lines using this RNAi system. We observed that TRMPV-PDCD10 cells treated with doxycycline showed reduced levels of PDCD10 (siPDCD10) when compared to controls (TRMPV-PDCD10 no doxycycline cells) and mimicked the transcriptomic changes relevant to CCM disease, similar to those observed in cultured primary mouse BEC(*19, 29*) (Supplemental Fig. 5). Notably, recent reports indicate that inflammation and hypoxia signaling are critical in the chronicity of CCM disease(*45*). Therefore, we investigated how a CCM-like environment(*45, 47*) influences changes in chromatin accessibility and gene expression in culture. This environment simulates a chronic inflammatory condition by utilizing TNFα (10 ng/ml) as an inflammatory stimulus and a prolyl hydroxylase inhibitor, dimethyloxalylglycine (500 mM,DMOG), as a hypoxia stimulus (Fig. 6 and Supplemental Fig. 5). We conducted bulk RNA-seq and ATAC-seq on parallel samples under the same experimental conditions (Fig. 6A-6D). We found 16,661 accessible chromatin sites and 2,077 genes upregulated in siPDCD10 BEC when exposed to a CCM-like environment. Moreover, we observed a significant correlation between siPDCD10 in a CCM-like environment, at both gene expression level (Pearson correlation, Pearson correlation, R = 0.48, *p* < 2.2x10^16^) and in chromatin accessibility (Pearson correlation, Pearson correlation, R = 0.26, *p* < 2.2x10^16^) to that identified in CCM endothelial cells derived from freshly isolated BEC in *Pdcd10^BECKO^* mice, as assessed by single-nucleus multi-omic analysis (Fig. 6A, 6B). Notably, we observed that a CCM-like environment itself led to increased accessibility at 18,809 sites and upregulation of 1,920 genes in TRMPV-*PDCD10* BEC without doxycycline (Controls) (Supplemental Fig. 5). We observed that the analysis of transcription factor motif enrichment in the predicted putative enhancer of BEC with siPDCD10, in a CCM-like environment, showed a high correlation (Pearson correlation, R = 0.83, *p* < 2.2x10^16)^ with the transcription factor motif enrichment found in subtype large veins of *Pdcd10^BECKO^* mice. To investigate the regulatory pathways related to chronic inflammation in brain endothelial cells, we performed an integrative analysis of our bulk RNA-seq and ATAC-seq data. By analyzing the correlation between the activity of brain endothelial cCREs in promoters associated with gene expression located within 500 kb, we identified 23,795 potential enhancers associated with 6,939 genes in human BEC. For this analysis, we used all upregulated genes in siPDCD10 under a CCM-like environment as our input. Furthermore, we performed a KEGG pathway enrichment analysis to identify the common regulatory systems influenced by these accessible putative enhancers and gene expression, which may drive the persistent activation of the BEC during neuroinflammation (Fig. 6D). The analysis revealed significant enrichment for terms previously associated with CCM exacerbation in human and animal models, including PI3K-Akt, Rap1, NLRP3, inflammation, and hypoxia pathways (Fig. 5D). We also observed that a CCM-like environment, even in the absence of CCM disease (TRMPV-PDCD10 cells), is sufficient to reprogram BEC toward a persistently activated state (Fig. 5D).

**Figure 6.**
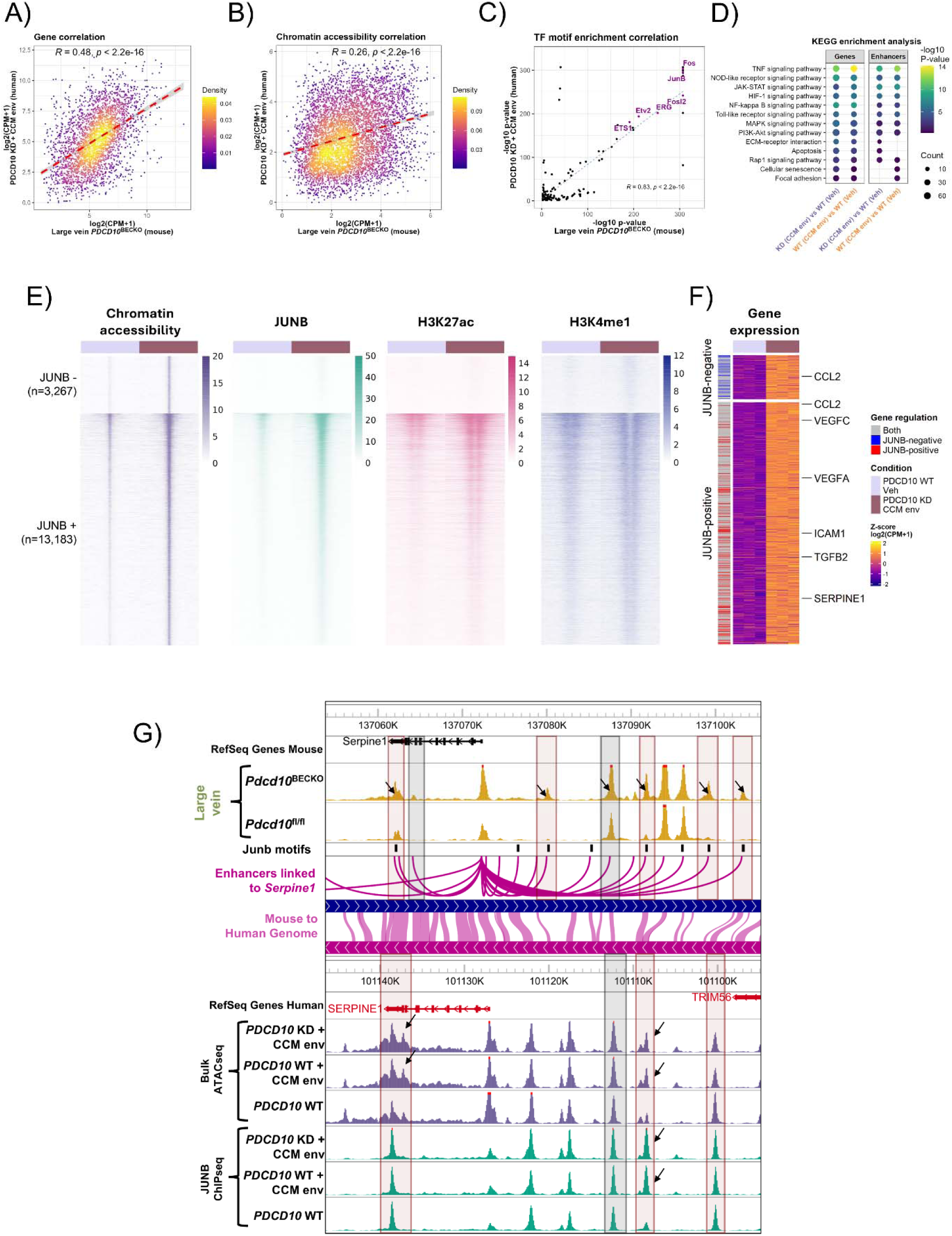
JUNB contributes to a persistent activated brain endothelial cell state. **A)** Scatter plot showing Pearson correlation of orthologous DEG between mouse *Pdcd10*^BECKO^ large vein BECs and human *PDCD10*-Knockdown (KD) cells under a CCM-like environment (500 µM DMOG and 10 ng/mL TNFα). Mouse values (x-axis) represent log_2_(CPM+1) from *Pdcd10*^BECKO^ large vein cells. Human values (y-axis) represent log_2_(CPM+1) from PDCD10 KD + CCM environment. The color scale indicates local 2D point density (yellow: High-density; purple: low-density). Pearson correlation is R = 0.49, *p* < 2.2^-16^. **B)** Correlation of orthologous chromatin accessibility shown as log₂(CPM+1) between *Pdcd10*^BECKO^ large vein BECs and human *PDCD10*-KD under CCM environment. Color scale as in **(A).** Pearson correlation is R = 0.26, *p* < 2.2^-16^. **C)**Scatter plot comparing TF motif enrichment -log_10_(p-value) between *Pdcd10*^BECKO^ large vein BECs and human *PDCD10*-KD under CCM environment. Each point represents a TF motif. Dashed blue line represents the regression line showing a positive correlation. Several conserved TFs, including JunB, Fos, Ets1, and ERG, are highlighted in magenta. Pearson correlation R = 0.53, p < 2.2e–16.**D)** KEGG pathway enrichment analysis of upregulated genes and more accessible putative enhancers in *PDCD10*-KD cells ± CCM environment compared to *PDCD10*-WT. Dot size represents the number of genes per term; color indicates -log10 *p*-value. Veh: Vehicle; CCM env: CCM environment. **E)** Heatmaps showing chromatin accessibility, JUNB binding, H3K27ac, and H3K4me1 levels at conserved cis-regulatory elements (cCREs) differentially accessible in PDCD10-KD + CCM env. Each row represents the same genomic region across all heatmaps. Regions were grouped into JUNB-positive (bottom) and JUNB-negative (top) categories based on JUNB binding. Color scale represents RPKM. **F)** Gene expression heatmap (Z-score of RNA log_2_ [CPM + 1]) for genes linked to region classified as putative enhancers from cCREs in **(D)**. Left annotation indicates whether the gene is associated with JUNB-dependent (red), JUNB-independent (blue), or both (gray) putative enhancer regions. **G)** WashU Comparative Epigenome Browser snapshots of *SERPINE1* locus. Tracks show chromatin accessibility, putative enhancer-gene linkage, and JUNB motifs in mouse large veins (*Pdcd10*^BECKO^ and *Pdcd10*^fl/fl^), alongside aligned human genomic region (hg38) displaying chromatin accessibility and JUNB binding in *PDCD10* WT under CCM-like environment, *PDCD10*-KD under CCM-like environment, and *PDCD10*-WT. Shared chromatin-accessible regions between mouse and human are highlighted in gray; shared regions containing JUNB motifs are highlighted in orange.

**Figure 7.**
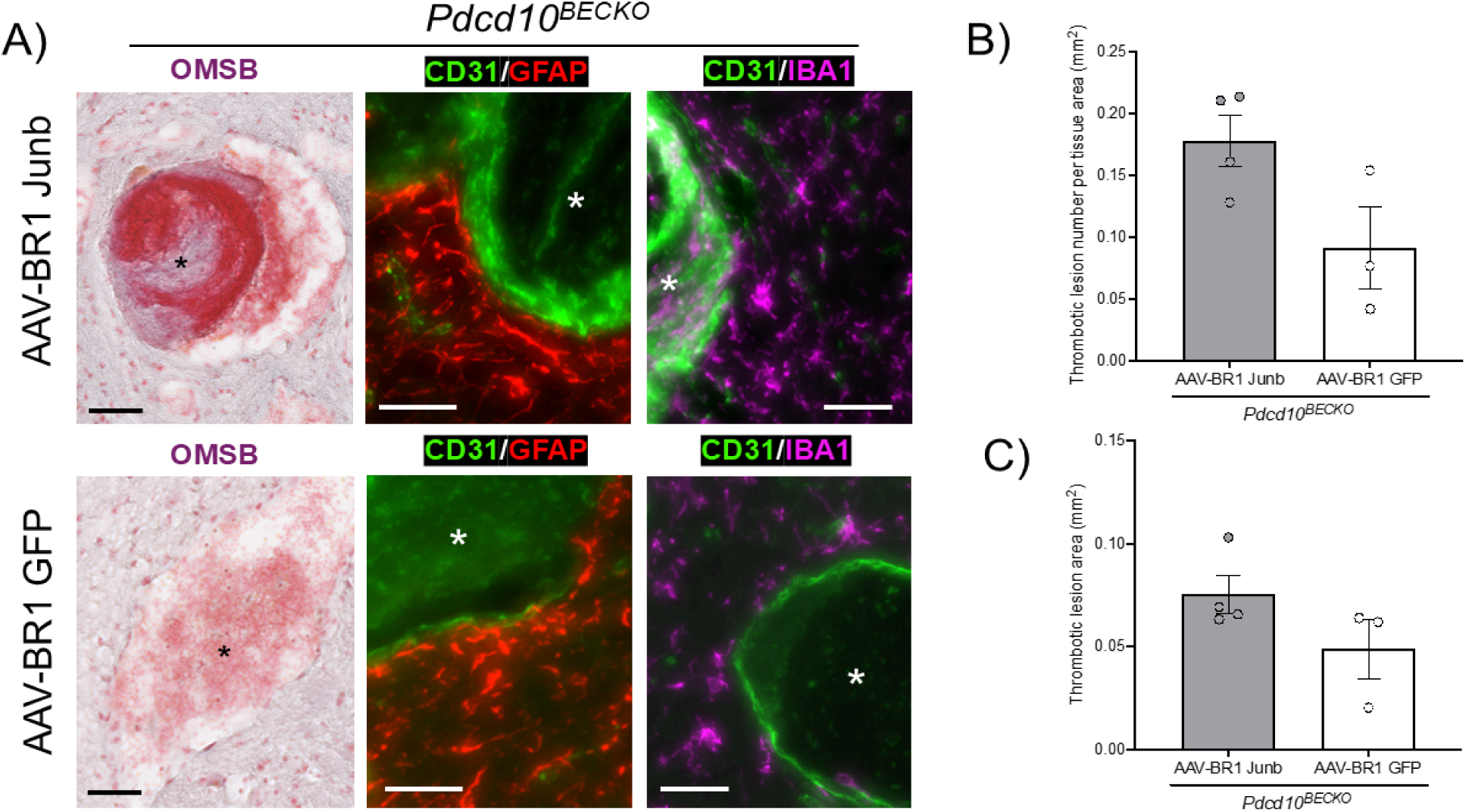
Ectopic expression of JUNB increases immunothrombosis in *Pdcd10^BECKO^* mice. **A)** Brain thrombi sections stained by OMSB, and immunofluorescence for GFAP+ astrocytes (red) and IBA1+ microglia and immune cells (magenta) in P60 *Pdcd10^BECKO^* mice treated with AAV-BR1 JUNB or AAV-BR1 GFP (control). Brain endothelium visualized with CD31 (green). **B)** Quantification of thrombus number per mm^2^ of brain section in *Pdcd10^BECKO^* mice treated with AAV-BR1 JUNB versus AAV-BR1 GFP. **C)** Quantification of thrombus area in *Pdcd10^BECKO^* mice treated with AAV-BR1 JUNB or AAV-BR1 GFP. All data are mean ± SEM, AAV-BR1 Junb treated n= 4; Control AAV-BR1 GFP treated n = 3. Black scale bars: 100 µm; white scale bars = 50 µm. Stars indicate lumen of CCM lesions.

Notably, the in vitro model of human CCM, combined with a multi-omic integration analysis of BEC and histological examinations of human and mouse brains affected by CCM, strongly suggests that the AP1 transcription factor JUNB may be associated with the reprogramming of brain endothelium towards a persistent activated cells state during neuroinflammation (Supplemental Fig. 5). To assess the potential role of JUNB in endothelial cell reprogramming during neuroinflammation, we performed chromatin immunoprecipitation followed by sequencing (ChIP-seq) targeting JUNB, the active enhancer histone modification H3K27ac, and the poised/active enhancer histone modification H3K4me1(*67-69*) (Fig. 6D-E). We observed a significant increase in JUNB-bound cCREs that became accessible under CCM conditions during chronic inflammation (in siPDCD10 and CCM-like environments), and had elevated H3K27ac levels. In contrast, we observed that cCREs without JUNB binding were devoid of H3K27ac. Notably, the H3K27ac-gained cCREs are marked with H3K4me1, suggesting a poised enhancer chromatin state at these cCREs prior to JUNB recruitment and nucleosome remodeling (Fig. 6E). These results indicate that JUNB binds to cCREs related to heightened endothelial inflammation, thrombosis, and dysfunction, as well as other active transcription sites identified by H3K27ac (Fig. 6F, Supplemental Fig. 5). Additionally, we found that the AP-1 inhibitor T5224 (40 µM) significantly reduces the recruitment of JUNB at inflammatory loci and decreases gene expression associated with an inflammatory state (Supplemental Fig. 5).

We next conducted a comparative analysis using the WashU Comparative Epigenome Browser(*119*), which revealed that several putative enhancers predicted in mice are conserved in humans. We identified direct regulation of PAI-1 (*SERPINE1*) expression through putative enhancers that are conserved between the human and mouse genomes (Fig. 6G). Our analysis further predicts that the AP-1 transcription factor JUNB act primarily as an activator in regulating PAI-1 expression in brain endothelial cells during chronic inflammation, as JUNB motifs were primarily found in cCREs and putative enhancers (Fig. 6G). These findings suggest that increased activity of the brain endothelial JUNB transcription factor, along with other members of the AP-1 family, may be linked to changes in chromatin accessibility during chronic CCM disease, which is associated with chronic inflammation. These changes contribute to shifts in gene expression programs that keep endothelial cells persistently activated due to chromatin accessibility at inflammatory loci. Importantly, the results indicate that this persistent activation of endothelial cells caused by inflammation and hypoxia occurs independently of the loss of CCM in brain endothelial cells.

### Elevation of brain endothelial AP-1 transcription factor JUNB exacerbates CCM disease

We next aimed to determine the causal links between the elevation of brain endothelial JUNB and CCM pathogenesis. Our approach involved utilizing a brain endothelial-targeted viral vector, specifically the adeno-associated virus (AAV)-BR1, to mediate the ectopic expression of JUNB in the brain endothelium specifically(*120*). In this experiment, we injected AAV-BR1-JUNB (1x10^11 vg/mouse) or control AAV-BR1-GFP retro-orbitally at P40 and at P45 in *Pdcd10^BECKO^* and *Pdcd10^fl/fl^* mice and assessed their brains at P60. Histological analysis, utilizing orcein and Martius Scarlet blue (OMSB)(*47*) staining, demonstrated that the ectopic expression of endothelial JUNB leads to increased lesion burden and thrombosis in *Pdcd10^BECKO^* treated with AAV-BR1-JUNB compared to *Pdcd10^BECKO^*treated with AAV-BR1-GFP. Moreover, immunofluorescent analysis further showed a more prominent presence of inflammation in *Pdcd10^BECKO^* treated with AAV-BR1-JUNB compared to *Pdcd10^BECKO^* treated with AAV-BR1-GFP. However, we found that the ectopic expression of endothelial JUNB did not significantly impact the brain vasculature in non-CCM mice (*Pdcd10^fl/fl^* treated with AAV-BR1-JUNB). These results suggest that increasing levels of brain endothelial JUNB alone are insufficient to reprogram brain endothelial cells. However, the elevation of brain endothelial JUNB may aid the reprogramming of these cells under specific conditions, such as inflammation and hypoxia.

## Discussion

In this study, we simultaneously profiled the brain endothelial transcriptome and chromatin accessibility using 10x Genomics snRNA-seq and snATAC-seq, revealing that epigenetic regulation plays a significant role in influencing the identity and function of brain endothelial subtypes within the arteriovenous axis. Moreover, through the integration of multi-omics analyses, we observed extensive epigenomic reprogramming of brain endothelial cell subtypes in a chronic model of CCM disease, characterized by a profound neuroinflammatory gene expression profile. We identified that the brain endothelial JUNB acted as an activator that regulates inflammatory loci and maintains a persistent activated cell state in chronic neuroinflammatory CCM animal models. Additionally, we uncovered both trans- and cis-regulatory factors in brain endothelial cells that are conserved between mice and humans, contributing to the progression of chronic CCM disease and susceptibility to inflammation and thrombosis.

The identification of specific subtypes of brain endothelial cells that express particular TFs and transporters, as well as differences in the expression of arterial and venous marker genes, has enhanced our understanding of the functional diversity within these cells. This knowledge has also provided a clearer understanding of their specialized roles along the arteriovenous axis in the brain(*91-98, 105*). In our study, we enriched brain endothelial cells from brain tissue, which are typically scarce in single-cell analyses. This approach enables us to collect comprehensive data on various subtypes of brain endothelial cells in both healthy mice and those with CCM disease. Our findings reveal that the identity, function, and gene expression patterns of each brain endothelial subtype along the arteriovenous axis are significantly influenced by conserved cCREs and putative enhancers in healthy mice. The identification of brain endothelial cCREs provide an opportunity to further understand the specific regulatory mechanisms by which brain endothelial TFs and chromatin-remodeling proteins interact with promoters, enhancers, silencers, insulators, and other known regulatory sequences(*101, 121, 122*) to carry out their specialized roles in the central nervous system. Our in-silico analysis, conducted through transcription motif enrichment and gene expression, suggested that the identity and function of brain endothelial cell subtypes are highly regulated by a network of transcriptional regulation. This network comprises a core group of TFs, which includes AP-1, SOX, GATA, and FOX, that help maintain the identity of brain endothelial subtypes and support their functional activity. Other families of TFs, such as ETS, KLF, and NF-kB, are equally important and likely help preserve the overall identity of endothelial cells(*93, 123*). Additional TFs, such as MEF2, NRF2, and STAT6, may regulate crucial vascular functions tailored to each specific brain endothelial subtype. Consistent with earlier reports, transcription factor motifs related to the SOX, TCF, and FOX transcription factor families have been previously noted to be more abundant in brain endothelial cells compared to peripheral endothelial cells(*93, 124*). For instance, SOX17 has been reported to regulate Wnt/β-catenin signaling, an important module involved in the induction and maintenance of the blood-brain barrier (BBB)(*125*). Our analysis revealed that the epigenome influences the expression of a group of genes crucial to endothelial processes, including angiogenesis, inflammatory response, and cellular metabolism, which are essential for the functions of brain endothelial cells. However, we did not observe a significant differential correlation among each brain endothelial subtype regarding the epigenome and the expression of genes involved in the BBB(*126*) through the arteriovenous axis in healthy mouse brains. Future studies should focus on the role of brain endothelial TF networks and their impact on the epigenome, which determines the specific programs and functions of various brain endothelial cell subtypes.

Although inflammation is known to contribute to the exacerbation of CCM disease(*45, 47*), the specific molecular mechanisms that lead to brain endothelial cell dysfunction, resulting in an increased risk of thrombosis, ongoing inflammation, and endothelial dysfunction, are poorly understood. This study demonstrates significant epigenomic reprogramming of brain endothelial cell subtypes in a neuroinflammatory CCM disease model. The study reveals how different subtypes of brain endothelial cells respond to chronic neuroinflammation and provides a comprehensive atlas of cCREs within the arteriovenous axis. This was achieved by profiling accessible chromatin and gene expression in individual nuclei. We observed significant changes in capillaries, capillary veins, and large veins, which show increased chromatin accessibility at gene promoters and distal regions associated with chronic inflammation and thrombotic responses. These changes are correlated with alterations in gene expression. These included genes involved in coagulation; EPCR (*Procr*), TM (*Thbd*), PAI-1 (*Serpine1*), TSP1 (*Thbs1*), VWF, and tPA (*Plat*), genes involved in antigen presentation; CD74, H2-A, H2-E, H2-DM, Chemokines; CX3CL1, ACRK, Cytokines and inflammatory response; IL6, HIF1a, FN1, TGFB1(*17, 45-47, 49*). However, the changes were less prominent in the subtypes of brain endothelial cells classified as artery. These findings are consistent with previous studies, which indicate that CCMs primarily affect veins and capillaries more than arteries(*18*). Future studies should investigate whether veins and capillaries are more susceptible to a stronger neuroinflammatory response due to their unique epigenetic profile. This could partially explain why veins are more likely to activate gene regulatory programs that promote neuroinflammation more readily than arteries. Understanding this mechanism will provide deeper insights into the pathogenesis of neurovascular inflammation and thrombosis.

A multi-omic integration analysis of brain endothelial cells, along with histological studies in both human and mouse brains affected by CCM, as well as an in vitro model of human CCM, identified the AP1 transcription factor JUNB as being highly expressed in thrombotic CCM lesions and correlated with reprogramming of the brain endothelium in chronic inflammation. AP-1 transcription factors plays a significant role in the maintenance of cancer(*60*), chronic inflammation(*127-129*), and coronary artery disease(*62*). Previous studies have shown that external signals modulate JUNB expression and act as an activator in cis-regulatory regions, which helps promote the differentiation of Th17 cells in chronic inflammatory diseases(*71, 72*). Furthermore, JUNB levels have been observed to increase following ICH and are associated with long-term neurological deficits(*77*). Here we show that JUNB in brain endothelial cells functions as an activator and binds to regulatory elements that exhibit increased levels of H3K27 acetylation (H3K27ac), a marker of active enhancers. This indicates a role for JUNB in inducing H3K27ac(*67-69*) at thousands of loci, including those associated with persistent activation of brain endothelial cells. These changes in gene expression increase the susceptibility of endothelial cells to thrombosis, inflammation, and dysfunction. Therefore, our research indicates that a multi-omic analysis of brain endothelial cells can facilitate the identification of various subtypes of these cells and their responses to chronic neuroinflammation. It also reveals the distinct endothelial cell states and molecular mechanisms that lead to a persistently activated endothelial cell state, which exhibits a high tendency for thrombosis, inflammation, and dysfunction. These insights could lead to new therapeutic approaches aimed at preventing neuroinflammatory diseases, thrombosis, and facilitating recovery.

## Authors’ contributions

H.G-G., designed and performed the experiments, analyzed and interpreted the data, generated the figures, and wrote the manuscript. E.F-A. designed and prepared all illustrations, designed, performed, analyzed histologically, interpreted the data, and generated the figures from experiments in conditional knockout mice. C.B., L.Z., helped with animal experiments and interpretation of results. E.H. designed and performed the experiments and analyzed the data of ChIP-seq experiments in cultured cells. H.S.I., performed 10x multiome experiments. J.K., J.T., J.S., provided critical reagents and interpretation of results and helped to review the manuscript. N.R.Z., & M.A.L-R. conceived the project, designed the overall study, analyzed and interpreted data, and wrote the manuscript.

## Acknowledgments

The authors thank Mark H. Ginsberg for helpful discussion; Brina Nguyen, Zaida Mizushima, and Nyle Connolly for technical assistance. *Slco1c1-CreERT2 mice* were the generous gift of Markus Schwaninger (University of Lu beck); *Pdcd10^fl/fl^*mice were the generous gift of Wang Min (Yale University).

## Figures

**Supplemental Figure 1.**
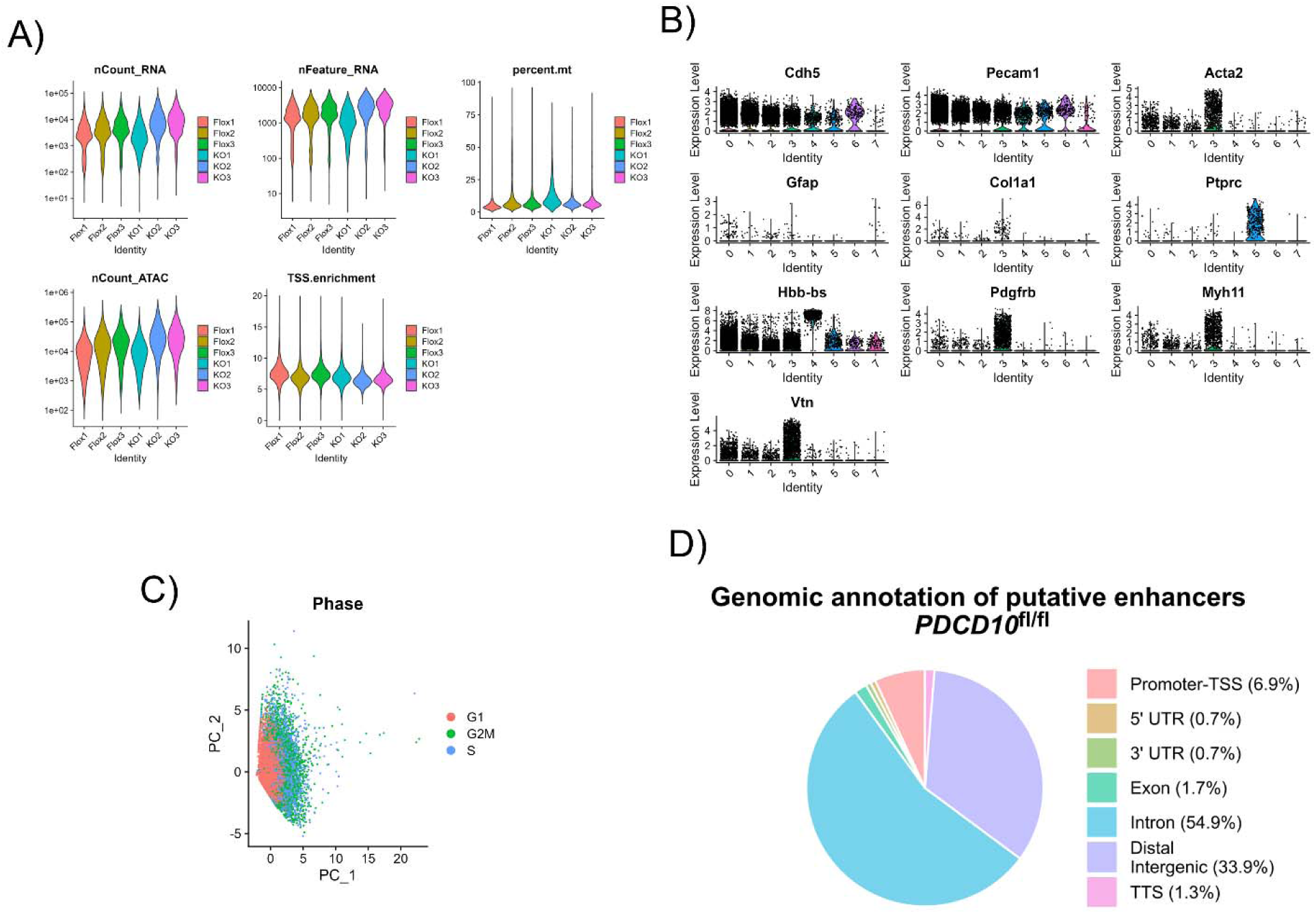
Single nuclei RNAseq/ATACseq quality control. **A)** Violin plots illustrating absolute RNA UMI count (nCount_RNA), RNA characteristic number (nFeature_RNA), percent of mitochondria genes (percent.mt), absolute ATAC UMI count (nCount_ATAC), and TSS enrichment (TSS.enrichment) before quality control filtering. *Pdc10^fl/fl^* (Flox) and *Pdcd10^BECKO^* (KO) number of biological replicates is indicated. **B)** Violin plots showing markers for pericytes (*Pdgfrb, Vtn, Myh11*), blood cells (*Hbb-bs*), and immune cells (*Ptprc*). These clusters were removed for downstream analysis. **C)** PCA plot of single cells colored by cell cycle phase. Cells are colored by assigned phase: G1 (red), S (blue), and G2/M (green). **D)** Genomic annotation of putative enhancers found in *Pdcd10*^fl/fl^ BEC subtypes.

**Supplemental Figure 2.**
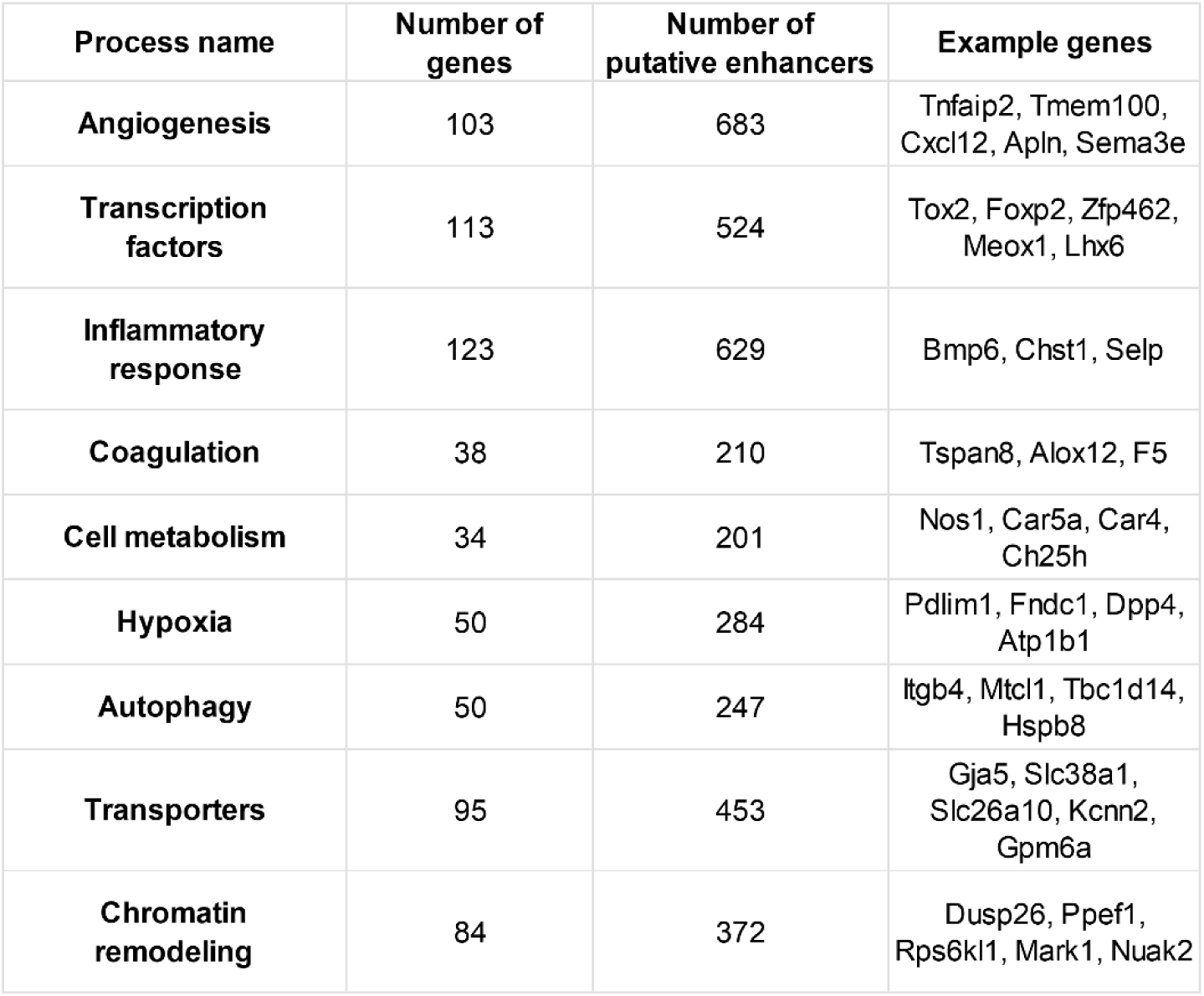
Summary of genes and putative enhancers by biological process in brain endothelial cells. The table lists selected biological processes, the number of genes and associated putative enhancers, and representative example genes for each process.

**Supplemental Figure 3.**
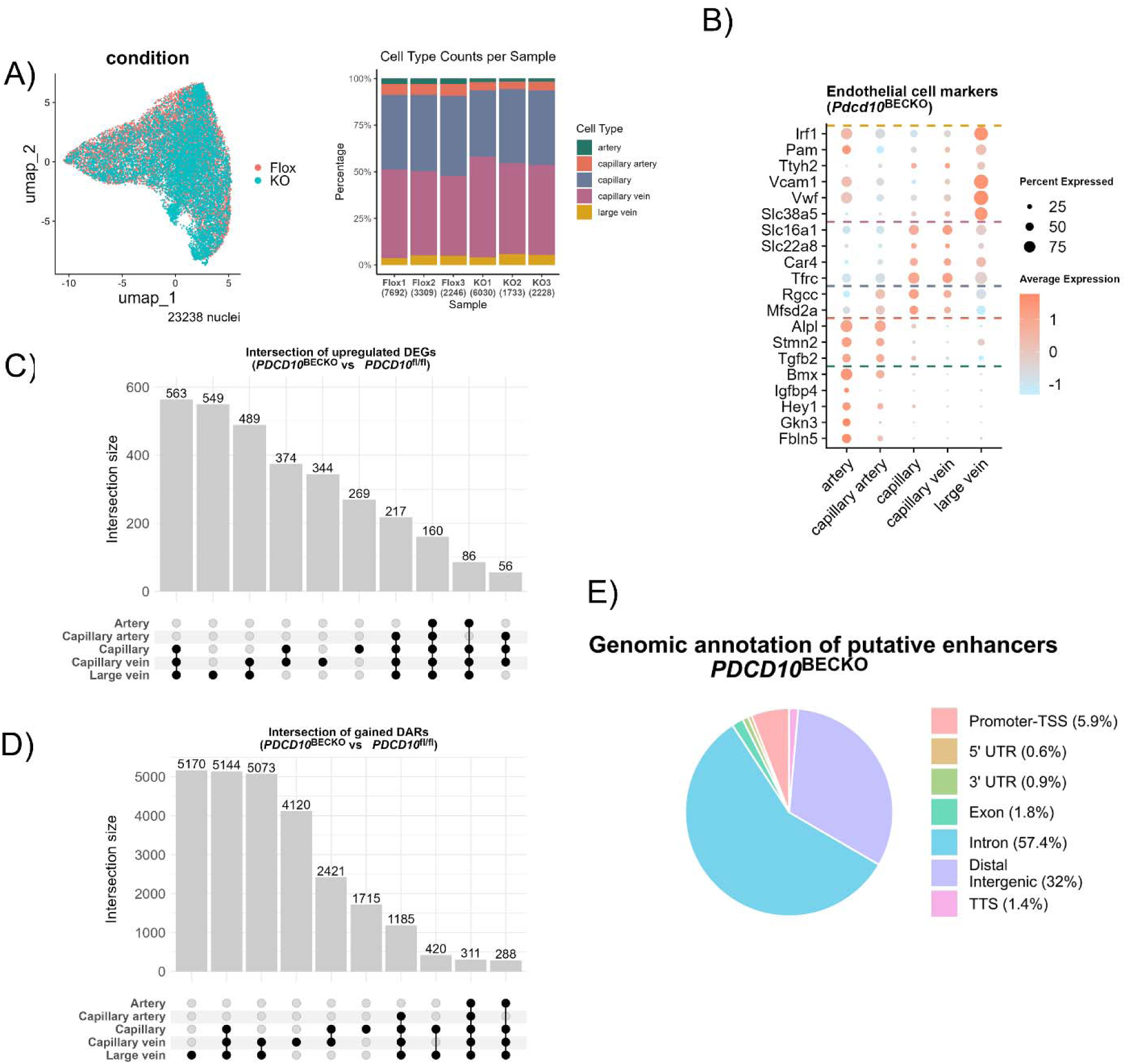
Multi-omic analysis of brain endothelial cells in *Pdcd10*^BECKO^ mice. **A)** Right: uniform manifold approximation and projection (UMAP) showing overlap of *Pdcd10*^BECKO^ and *Pdcd10*^fl/fl^ mouse. Left: distribution of BEC subtype (y-axis) across *Pdcd10*^fl/fl^ and *Pdcd10*^BECKO^ mouse replicates. Bar labels indicate the total number of nuclei per replicate. Colors correspond to the five BEC subtypes. **B)** Dot plot shows the average expression levels of BEC subtype markers and the percentage of expressing cells in *Pdcd10*^BECKO^. Dashed lines separate the five different BEC subtypes. **C)** Upset plots showing the intersection of genes along the arteriovenous axis. **D)** Upset plots showing the intersection of putative enhancers along the arteriovenous axis. **E)** Genomic annotation of putative enhancers found in *Pdcd10*^BECKO^ BEC subtypes.

**Supplemental Figure 4.**
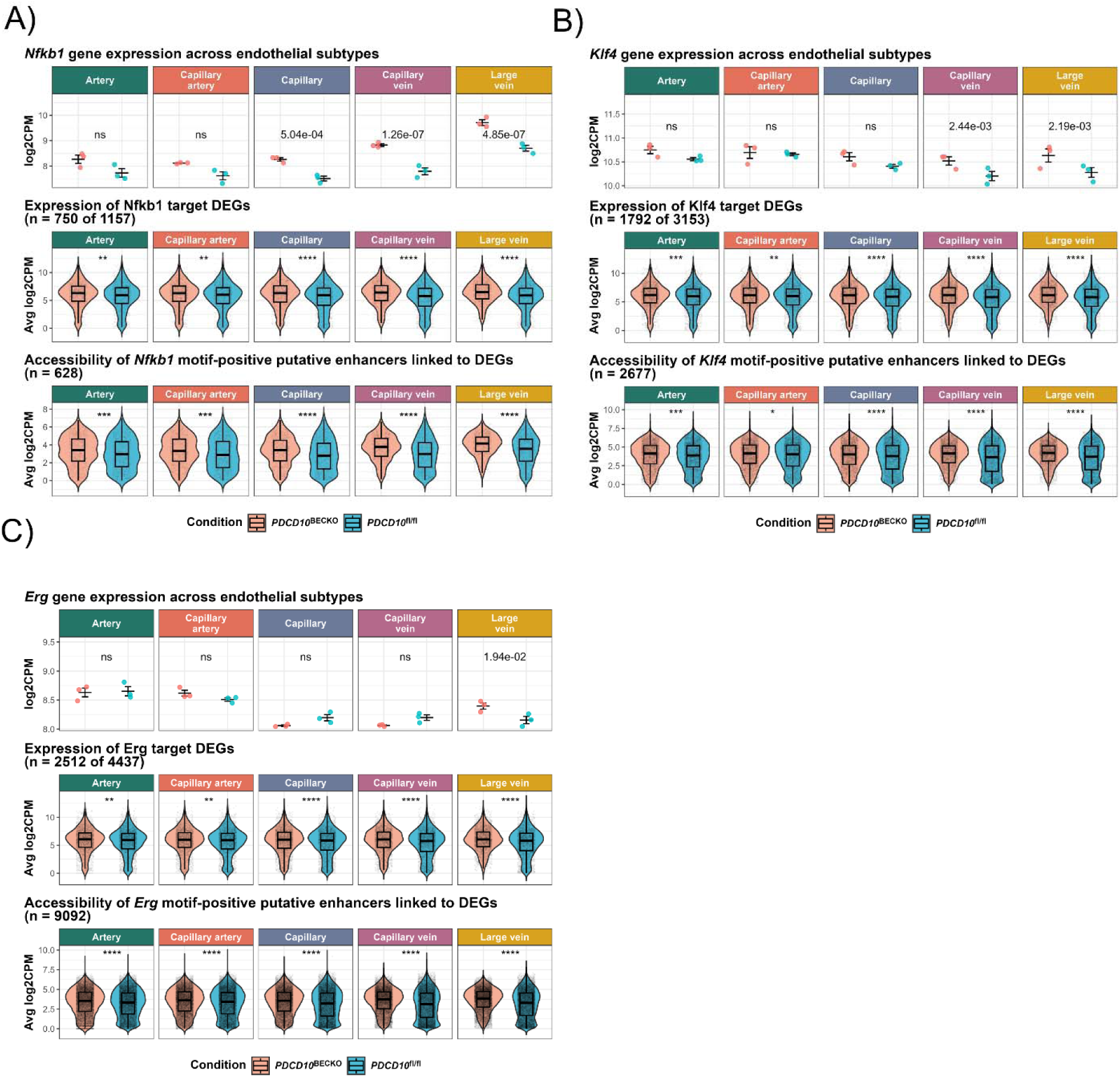
Expression and regulatory activity of Nfkb1, Klf4, and Erg across brain endothelial subtypes. A-C) Transcription-factor expression and enhancer activity of Nfkb1, Klf4, and Erg across BEC subtypes. Top: Gene expression in *Pdcd10*^BECKO^ (pink dots) and *Pdcd10*^fl/fl^ (blue dots) across endothelial subtypes. Each point represents a biological replicate. FDR values were calculated using edgeR’s *glmLRT* function; ns, not significant. **Middle:** Expression of differentially expressed genes (DEGs) regulated by the respective transcription factor across BEC subtypes. Each point represents one DEG. **Bottom:** Chromatin accessibility of motif-containing putative enhancers linked to DEGs, per BEC subtype. Each point represents an individual putative enhancer. For the **middle** and **bottom** panels, *P*-values from Wilcoxon rank-sum tests are shown. For the **top** panel, error bars represent mean ± SEM. Subtype-specific colors indicate the endothelial subtype.

**Supplemental Figure 5.**
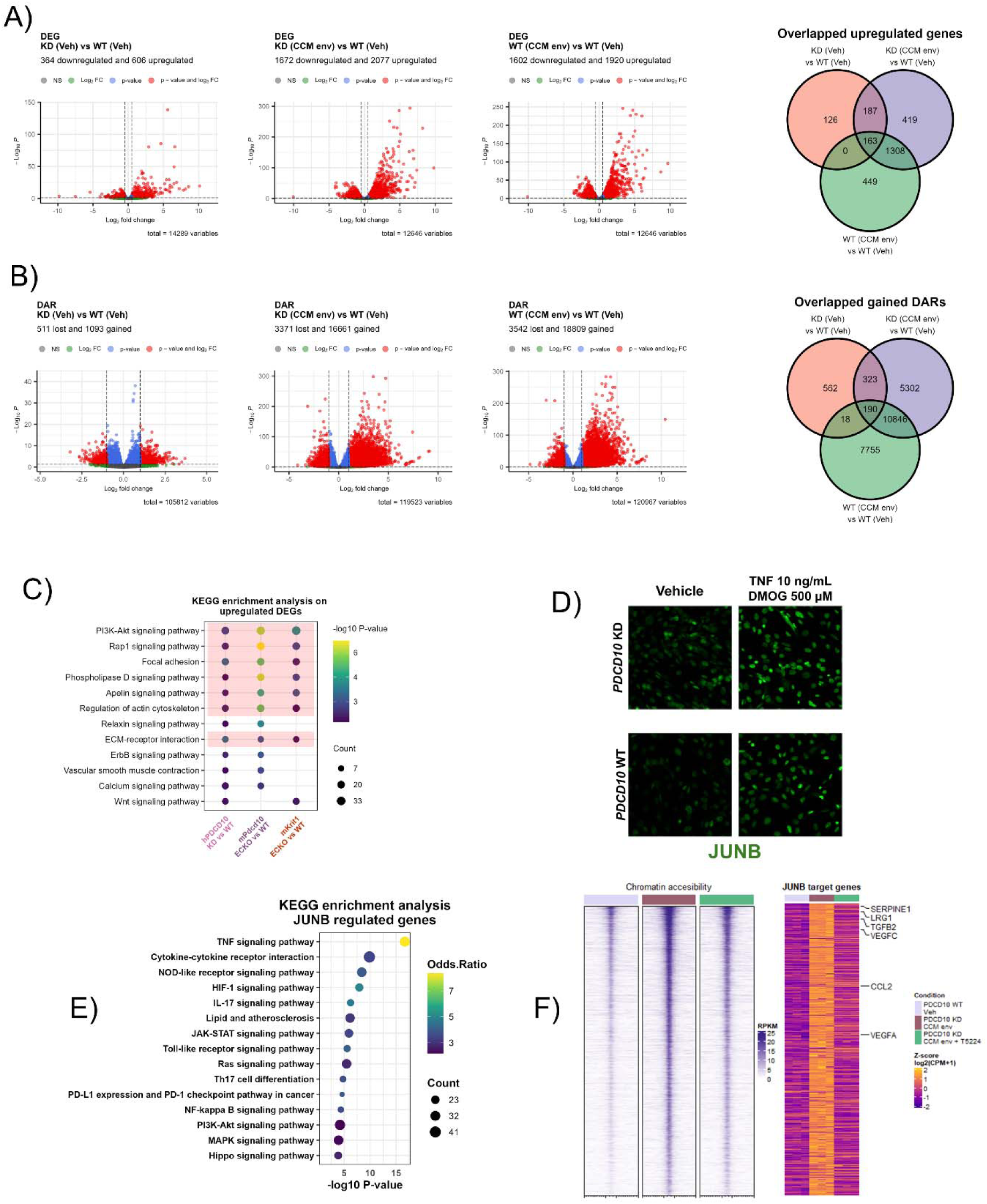
JUNB contributes to a persistent activated brain endothelial cell state in human brain endothelial cells. A) Volcano plots showing differentially expressed genes (DEGs) from PDCD10 KD, and PDCD10 KD and WT under CCM-like environment vehicle. Right: Venn diagram showing overlap of upregulated DEGs across conditions. B) Volcano plots of differentially accessible regions (DARs) in ATAC-seq from PDCD10 KD, and PDCD10 KD and WT under CCM-like environment vehicle. Right: Venn diagram showing overlap of accessible regions across conditions. C) KEGG pathway enrichment analysis of DEGs upregulated under CCM conditions compared with previous mouse primary endothelial cell Knock-out models of *Krit1* and *Pdcd10*. D) JUNB immunostaining in PDCD10 KD and WT BECs under vehicle or CCM-like environment after 6 hours, showing JUNB induction. F) KEGG enrichment analysis of JUNB-regulated genes, defined by JUNB motif-containing putative enhancers linked to DEGs. F) Heatmaps showing chromatin accessibility at JUNB-regulated regions in PDCD10 WT (light purple), PDCD10 KD under CCM environment (brown) and PDCD10 KD under CCM environment and AP-1 inhibitor (green). Expression of JUNB target genes (right heatmap) from the same conditions.

## Reference

1. Z. Morris et al., Incidental findings on brain magnetic resonance imaging: systematic review and meta-analysis. BMJ 339, b3016 (2009).

2. S. Spiegler et al., High mutation detection rates in cerebral cavernous malformation upon stringent inclusion criteria: one-third of probands are minors. Mol Genet Genomic Med 2, 176–185 (2014).

3. S. Weinsheimer et al., Intracranial Hemorrhage Rate and Lesion Burden in Patients With Familial Cerebral Cavernous Malformation. J Am Heart Assoc 12, e027572 (2023).

4. L. Morrison, A. Akers, in GeneReviews((R)), M. P. Adam et al., Eds. (Seattle (WA), 1993).

5. S. Spiegler, M. Rath, C. Paperlein, U. Felbor, Cerebral Cavernous Malformations: An Update on Prevalence, Molecular Genetic Analyses, and Genetic Counselling. Mol Syndromol 9, 60–69 (2018).

6. H. Choquet et al., Polymorphisms in inflammatory and immune response genes associated with cerebral cavernous malformation type 1 severity. Cerebrovasc Dis 38, 433–440 (2014).

7. T. Hong et al., Somatic MAP3K3 and PIK3CA mutations in sporadic cerebral and spinal cord cavernous malformations. Brain 144, 2648–2658 (2021).

8. M. Peyre et al., Somatic PIK3CA Mutations in Sporadic Cerebral Cavernous Malformations. N Engl J Med 385, 996–1004 (2021).

9. A. A. Ren et al., PIK3CA and CCM mutations fuel cavernomas through a cancer-like mechanism. Nature 594, 271–276 (2021).

10. A. Perrelli, S. F. Retta, Polymorphisms in genes related to oxidative stress and inflammation: Emerging links with the pathogenesis and severity of Cerebral Cavernous Malformation disease. Free Radic Biol Med 172, 403–417 (2021).

11. C. B. Josephson et al., Seizure risk from cavernous or arteriovenous malformations: prospective population-based study. Neurology 76, 1548–1554 (2011).

12. A. Fischer, J. Zalvide, E. Faurobert, C. Albiges-Rizo, E. Tournier-Lasserve, Cerebral cavernous malformations: from CCM genes to endothelial cell homeostasis. Trends Mol Med 19, 302–308 (2013).

13. B. N. Catherine Chinhchu Lai, Eduardo Frias-Anaya, Helios Gallego-Gutierrez, Marco Orecchioni, Victoria Herrera, Elan Ortiz, Hao Sun, Omar A. Mesarwi, Klaus Ley, Brendan Gongol & Miguel Alejandro Lopez-Ramirez, Neuroinflammation plays a critical role in cerebral cavernous malformation disease. Circ Res, (2022).

14. H. Choquet, L. Pawlikowska, M. T. Lawton, H. Kim, Genetics of cerebral cavernous malformations: current status and future prospects. J Neurosurg Sci 59, 211–220 (2015).

15. C. Denier et al., Genotype-phenotype correlations in cerebral cavernous malformations patients. Ann Neurol 60, 550–556 (2006).

16. A. G. Mikati et al., Vascular permeability in cerebral cavernous malformations. J Cereb Blood Flow Metab 35, 1632–1639 (2015).

17. M. A. Lopez-Ramirez et al., Cerebral cavernous malformations form an anticoagulant vascular domain in humans and mice. Blood 133, 193–204 (2019).

18. G. Boulday et al., Developmental timing of CCM2 loss influences cerebral cavernous malformations in mice. J Exp Med 208, 1835–1847 (2011).

19. M. A. Lopez-Ramirez et al., Thrombospondin1 (TSP1) replacement prevents cerebral cavernous malformations. J Exp Med 214, 3331–3346 (2017).

20. S. M. Stamatovic, N. Sladojevic, R. F. Keep, A. V. Andjelkovic, PDCD10 (CCM3) regulates brain endothelial barrier integrity in cerebral cavernous malformation type 3: role of CCM3-ERK1/2-cortactin cross-talk. Acta Neuropathol 130, 731–750 (2015).

21. A. Glading, J. Han, R. A. Stockton, M. H. Ginsberg, KRIT-1/CCM1 is a Rap1 effector that regulates endothelial cell cell junctions. J Cell Biol 179, 247–254 (2007).

22. K. J. Whitehead, N. W. Plummer, J. A. Adams, D. A. Marchuk, D. Y. Li, Ccm1 is required for arterial morphogenesis: implications for the etiology of human cavernous malformations. Development 131, 1437–1448 (2004).

23. P. V. DiStefano, J. M. Kuebel, I. H. Sarelius, A. J. Glading, KRIT1 protein depletion modifies endothelial cell behavior via increased vascular endothelial growth factor (VEGF) signaling. J Biol Chem 289, 33054–33065 (2014).

24. R. Girard et al., Plasma Biomarkers of Inflammation and Angiogenesis Predict Cerebral Cavernous Malformation Symptomatic Hemorrhage or Lesional Growth. Circ Res 122, 1716–1721 (2018).

25. J. Wustehube et al., Cerebral cavernous malformation protein CCM1 inhibits sprouting angiogenesis by activating DELTA-NOTCH signaling. Proc Natl Acad Sci U S A 107, 12640–12645 (2010).

26. L. Maddaluno et al., EndMT contributes to the onset and progression of cerebral cavernous malformations. Nature 498, 492–496 (2013).

27. L. Goitre et al., KRIT1 regulates the homeostasis of intracellular reactive oxygen species. PLoS One 5, e11786 (2010).

28. L. Goitre et al., KRIT1 loss of function causes a ROS-dependent upregulation of c-Jun. Free Radic Biol Med 68, 134–147 (2014).

29. J. Koskimaki et al., Comprehensive transcriptome analysis of cerebral cavernous malformation across multiple species and genotypes. JCI Insight 4, (2019).

30. H. J. Zhou et al., Erratum: Endothelial exocytosis of angiopoietin-2 resulting from CCM3 deficiency contributes to cerebral cavernous malformation. Nat Med 22, 1502 (2016).

31. A. C. Chan et al., Mutations in 2 distinct genetic pathways result in cerebral cavernous malformations in mice. J Clin Invest 121, 1871–1881 (2011).

32. D. A. McDonald et al., A novel mouse model of cerebral cavernous malformations based on the two-hit mutation hypothesis recapitulates the human disease. Hum Mol Genet 20, 211–222 (2011).

33. A. L. Akers, E. Johnson, G. K. Steinberg, J. M. Zabramski, D. A. Marchuk, Biallelic somatic and germline mutations in cerebral cavernous malformations (CCMs): evidence for a two-hit mechanism of CCM pathogenesis. Hum Mol Genet 18, 919–930 (2009).

34. K. J. Whitehead et al., The cerebral cavernous malformation signaling pathway promotes vascular integrity via Rho GTPases. Nat Med 15, 177–184 (2009).

35. D. A. McDonald et al., Fasudil decreases lesion burden in a murine model of cerebral cavernous malformation disease. Stroke 43, 571–574 (2012).

36. L. Bravi et al., Sulindac metabolites decrease cerebrovascular malformations in CCM3-knockout mice. Proc Natl Acad Sci U S A 112, 8421–8426 (2015).

37. C. C. Gibson et al., Strategy for identifying repurposed drugs for the treatment of cerebral cavernous malformation. Circulation 131, 289–299 (2015).

38. M. Reinhard et al., Propranolol stops progressive multiple cerebral cavernoma in an adult patient. J Neurol Sci 367, 15–17 (2016).

39. P. V. DiStefano, A. J. Glading, VEGF signalling enhances lesion burden in KRIT1 deficient mice. J Cell Mol Med 24, 632–639 (2020).

40. R. D’Angelo et al., Mutation analysis of CCM1, CCM2 and CCM3 genes in a cohort of Italian patients with cerebral cavernous malformation. Brain Pathol 21, 215–224 (2011).

41. G. Tanriover et al., Ultrastructural analysis of vascular features in cerebral cavernous malformations. Clinical neurology and neurosurgery 115, 438–444 (2013).

42. S. Y. Cheng, M. Nagane, H. S. Huang, W. K. Cavenee, Intracerebral tumor-associated hemorrhage caused by overexpression of the vascular endothelial growth factor isoforms VEGF121 and VEGF165 but not VEGF189. Proc Natl Acad Sci U S A 94, 12081–12087 (1997).

43. J. Josko, Cerebral angiogenesis and expression of VEGF after subarachnoid hemorrhage (SAH) in rats. Brain Res 981, 58–69 (2003).

44. I. A. Awad, S. P. Polster, Cavernous angiomas: deconstructing a neurosurgical disease. J Neurosurg 131, 1–13 (2019).

45. C. C. Lai et al., Neuroinflammation Plays a Critical Role in Cerebral Cavernous Malformation Disease. Circ Res 131, 909–925 (2022).

46. M. A. Lopez-Ramirez et al., Astrocytes propel neurovascular dysfunction during cerebral cavernous malformation lesion formation. J Clin Invest 131, (2021).

47. E. Frias-Anaya et al., Mild Hypoxia Accelerates Cerebral Cavernous Malformation Disease Through CX3CR1-CX3CL1 Signaling. Arterioscler Thromb Vasc Biol, (2024).

48. M. A. Globisch et al., Immunothrombosis and vascular heterogeneity in cerebral cavernous malformation. Blood, (2022).

49. A. C. Y. Yau et al., Inflammation and neutrophil extracellular traps in cerebral cavernous malformation. Cell Mol Life Sci 79, 206 (2022).

50. F. Lazzaroni et al., Circulating biomarkers in familial cerebral cavernous malformation. EBioMedicine 99, 104914 (2024).

51. S. Jauhiainen et al., Proteomics on human cerebral cavernous malformations reveals novel biomarkers in neurovascular dysfunction for the disease pathology. Biochim Biophys Acta Mol Basis Dis 1870, 167139 (2024).

52. A. T. Tang et al., Endothelial TLR4 and the microbiome drive cerebral cavernous malformations. Nature 545, 305–310 (2017).

53. S. M. Zuurbier et al., Long-term antithrombotic therapy and risk of intracranial haemorrhage from cerebral cavernous malformations: a population-based cohort study, systematic review, and meta-analysis. Lancet Neurol 18, 935–941 (2019).

54. D. A. Snellings et al., Cerebral Cavernous Malformation: From Mechanism to Therapy. Circ Res 129, 195–215 (2021).

55. Y. Li et al., Transcriptomic signatures of individual cell types in cerebral cavernous malformation. Cell Commun Signal 22, 23 (2024).

56. V. C. Pham et al., Epigenetic regulation by polycomb repressive complex 1 promotes cerebral cavernous malformations. EMBO Mol Med 16, 2827–2855 (2024).

57. C. M. Phillips, S. M. Stamatovic, R. F. Keep, A. V. Andjelkovic, Epigenetics and stroke: role of DNA methylation and effect of aging on blood-brain barrier recovery. Fluids Barriers CNS 20, 14 (2023).

58. Y. Zhao et al., Inhibition of endothelial histone deacetylase 2 shifts endothelial-mesenchymal transitions in cerebral arteriovenous malformation models. J Clin Invest 134, (2024).

59. M. Valentino et al., BMI1 Inhibition Improves Lesion Burden in Cerebral Cavernous Malformations. Circulation 150, 738–741 (2024).

60. Z. Wu, M. Nicoll, R. J. Ingham, AP-1 family transcription factors: a diverse family of proteins that regulate varied cellular activities in classical hodgkin lymphoma and ALK+ ALCL. Exp Hematol Oncol 10, 4 (2021).

61. M. Yukawa et al., AP-1 activity induced by co-stimulation is required for chromatin opening during T cell activation. J Exp Med 217, (2020).

62. Q. Zhao et al., TCF21 and AP-1 interact through epigenetic modifications to regulate coronary artery disease gene expression. Genome Med 11, 23 (2019).

63. S. C. Biddie et al., Transcription factor AP1 potentiates chromatin accessibility and glucocorticoid receptor binding. Mol Cell 43, 145–155 (2011).

64. Y. Zhang et al., Multi-omics computational analysis unveils the involvement of AP-1 and CTCF in hysteresis of chromatin states during macrophage polarization. Front Immunol 14, 1304778 (2023).

65. S. L. Klemm, Z. Shipony, W. J. Greenleaf, Chromatin accessibility and the regulatory epigenome. Nat Rev Genet 20, 207–220 (2019).

66. T. Ito et al., Identification of SWI.SNF complex subunit BAF60a as a determinant of the transactivation potential of Fos/Jun dimers. J Biol Chem 276, 2852–2857 (2001).

67. S. B. Larsen et al., Establishment, maintenance, and recall of inflammatory memory. Cell Stem Cell 28, 1758–1774 e1758 (2021).

68. R. Lin et al., H3K27ac mediated SS18/BAFs relocation regulates JUN induced pluripotent-somatic transition. Cell Biosci 12, 89 (2022).

69. J. Liao, J. Ho, M. Burns, E. C. Dykhuizen, D. C. Hargreaves, Collaboration between distinct SWI/SNF chromatin remodeling complexes directs enhancer selection and activation of macrophage inflammatory genes. Immunity 57, 1780–1795 e1786 (2024).

70. T. Vierbuchen et al., AP-1 Transcription Factors and the BAF Complex Mediate Signal-Dependent Enhancer Selection. Mol Cell 68, 1067–1082 e1012 (2017).

71. T. M. Carr, J. D. Wheaton, G. M. Houtz, M. Ciofani, JunB promotes Th17 cell identity and restrains alternative CD4(+) T-cell programs during inflammation. Nat Commun 8, 301 (2017).

72. M. Ciofani et al., A validated regulatory network for Th17 cell specification. Cell 151, 289–303 (2012).

73. F. J. Ren, X. Y. Cai, Y. Yao, G. Y. Fang, JunB: a paradigm for Jun family in immune response and cancer. Front Cell Infect Microbiol 13, 1222265 (2023).

74. Z. Y. Wang et al., Regulation of IL-10 gene expression in Th2 cells by Jun proteins. J Immunol 174, 2098–2105 (2005).

75. R. B. Ramos et al., Shock drives a STAT3 and JunB-mediated coordinated transcriptional and DNA methylation response in the endothelium. J Cell Sci 136, (2023).

76. P. Maity et al., Persistent JunB activation in fibroblasts disrupts stem cell niche interactions enforcing skin aging. Cell Rep 36, 109634 (2021).

77. R. Xiang et al., Spatiotemporal transcriptomic maps of mouse intracerebral hemorrhage at single-cell resolution. Neuron, (2025).

78. Y. Hao et al., Dictionary learning for integrative, multimodal and scalable single-cell analysis. Nat Biotechnol 42, 293–304 (2024).

79. C. S. McGinnis, L. M. Murrow, Z. J. Gartner, DoubletFinder: Doublet Detection in Single-Cell RNA Sequencing Data Using Artificial Nearest Neighbors. Cell Syst 8, 329–337 e324 (2019).

80. J. Kalucka et al., Single-Cell Transcriptome Atlas of Murine Endothelial Cells. Cell 180, 764–779 e720 (2020).

81. Y. E. Li et al., A comparative atlas of single-cell chromatin accessibility in the human brain. Science 382, eadf7044 (2023).

82. N. R. Zemke et al., Conserved and divergent gene regulatory programs of the mammalian neocortex. Nature 624, 390–402 (2023).

83. T. Stuart, A. Srivastava, S. Madad, C. A. Lareau, R. Satija, Single-cell chromatin state analysis with Signac. Nat Methods 18, 1333–1341 (2021).

84. F. Consortium et al., A promoter-level mammalian expression atlas. Nature 507, 462–470 (2014).

85. M. V. Kuleshov et al., Enrichr: a comprehensive gene set enrichment analysis web server 2016 update. Nucleic Acids Res 44, W90–97 (2016).

86. M. A. Lopez-Ramirez et al., Role of caspases in cytokine-induced barrier breakdown in human brain endothelial cells. J Immunol 189, 3130–3139 (2012).

87. M. A. Lopez-Ramirez et al., MicroRNA-155 negatively affects blood-brain barrier function during neuroinflammation. FASEB J 28, 2551–2565 (2014).

88. J. Zuber et al., Toolkit for evaluating genes required for proliferation and survival using tetracycline-regulated RNAi. Nat Biotechnol 29, 79–83 (2011).

89. M. R. Corces et al., An improved ATAC-seq protocol reduces background and enables interrogation of frozen tissues. Nat Methods 14, 959–962 (2017).

90. A. S. Hinrichs et al., The UCSC Genome Browser Database: update 2006. Nucleic Acids Res 34, D590–598 (2006).

91. N. R. Zemke et al., Author Correction: Conserved and divergent gene regulatory programs of the mammalian neocortex. Nature 625, E26 (2024).

92. K. Zhang, N. R. Zemke, E. J. Armand, B. Ren, A fast, scalable and versatile tool for analysis of single-cell omics data. Nat Methods 21, 217–227 (2024).

93. M. Yu et al., Integrative multi-omic profiling of adult mouse brain endothelial cells and potential implications in Alzheimer’s disease. Cell Rep 42, 113392 (2023).

94. E. E. Crouch, T. Joseph, E. Marsan, E. J. Huang, Disentangling brain vasculature in neurogenesis and neurodegeneration using single-cell transcriptomics. Trends Neurosci 46, 551–565 (2023).

95. A. Zeisel et al., Molecular Architecture of the Mouse Nervous System. Cell 174, 999–1014 e1022 (2018).

96. F. J. Garcia et al., Single-cell dissection of the human brain vasculature. Nature 603, 893–899 (2022).

97. T. Walchli et al., Single-cell atlas of the human brain vasculature across development, adulthood and disease. Nature 632, 603–613 (2024).

98. M. Vanlandewijck et al., Author Correction: A molecular atlas of cell types and zonation in the brain vasculature. Nature 560, E3 (2018).

99. J. Feng, T. Liu, B. Qin, Y. Zhang, X. S. Liu, Identifying ChIP-seq enrichment using MACS. Nat Protoc 7, 1728–1740 (2012).

100. S. Ma et al., Chromatin Potential Identified by Shared Single-Cell Profiling of RNA and Chromatin. Cell 183, 1103–1116 e1120 (2020).

101. S. Preissl, K. J. Gaulton, B. Ren, Characterizing cis-regulatory elements using single-cell epigenomics. Nat Rev Genet 24, 21–43 (2023).

102. S. Zu et al., Single-cell analysis of chromatin accessibility in the adult mouse brain. Nature 624, 378–389 (2023).

103. T. Stuart, A. Srivastava, S. Madad, C. A. Lareau, R. Satija, Author Correction: Single-cell chromatin state analysis with Signac. Nat Methods 19, 257 (2022).

104. S. Heinz et al., Simple combinations of lineage-determining transcription factors prime cis-regulatory elements required for macrophage and B cell identities. Mol Cell 38, 576–589 (2010).

105. L. Zhao et al., Pharmacologically reversible zonation-dependent endothelial cell transcriptomic changes with neurodegenerative disease associations in the aged brain. Nat Commun 11, 4413 (2020).

106. M. Vanlandewijck et al., A molecular atlas of cell types and zonation in the brain vasculature. Nature 554, 475–480 (2018).

107. C. Maderna, F. Pisati, C. Tripodo, E. Dejana, M. Malinverno, A murine model of cerebral cavernous malformations with acute hemorrhage. iScience 25, 103943 (2022).

108. E. A. Winkler et al., A single-cell atlas of the normal and malformed human brain vasculature. Science 375, eabi7377 (2022).

109. C. Shi et al., B-Cell Depletion Reduces the Maturation of Cerebral Cavernous Malformations in Murine Models. J Neuroimmune Pharmacol 11, 369–377 (2016).

110. M. Kanehisa, M. Furumichi, Y. Sato, Y. Matsuura, M. Ishiguro-Watanabe, KEGG: biological systems database as a model of the real world. Nucleic Acids Res 53, D672–D677 (2025).

111. E. Y. Chen et al., Enrichr: interactive and collaborative HTML5 gene list enrichment analysis tool. BMC Bioinformatics 14, 128 (2013).

112. Z. Xie et al., Gene Set Knowledge Discovery with Enrichr. Curr Protoc 1, e90 (2021).

113. C. D. Much et al., Inactivation of Cerebral Cavernous Malformation Genes Results in Accumulation of von Willebrand Factor and Redistribution of Weibel-Palade Bodies in Endothelial Cells. Front Mol Biosci 8, 622547 (2021).

114. M. F. Fontana et al., JUNB is a key transcriptional modulator of macrophage activation. J Immunol 194, 177–186 (2015).

115. P. Novoszel et al., The AP-1 transcription factors c-Jun and JunB are essential for CD8alpha conventional dendritic cell identity. Cell Death Differ 28, 2404–2420 (2021).

116. M. Renz et al., Regulation of beta1 integrin-Klf2-mediated angiogenesis by CCM proteins. Dev Cell 32, 181–190 (2015).

117. Z. Zhou et al., Corrigendum: Cerebral cavernous malformations arise from endothelial gain of MEKK3-KLF2/4 signalling. Nature 536, 488 (2016).

118. R. Cuttano et al., KLF4 is a key determinant in the development and progression of cerebral cavernous malformations. EMBO Mol Med 8, 6–24 (2016).

119. C. Seng et al., WashU Epigenome Browser update 2025. Nucleic Acids Res, (2025).

120. J. Korbelin et al., A brain microvasculature endothelial cell-specific viral vector with the potential to treat neurovascular and neurological diseases. EMBO Mol Med 8, 609–625 (2016).

121. V. Agarwal et al., Massively parallel characterization of transcriptional regulatory elements. Nature 639, 411–420 (2025).

122. K. Zhang et al., A single-cell atlas of chromatin accessibility in the human genome. Cell 184, 5985–6001 e5919 (2021).

123. M. F. Sabbagh et al., Transcriptional and epigenomic landscapes of CNS and non-CNS vascular endothelial cells. Elife 7, (2018).

124. M. Hupe et al., Gene expression profiles of brain endothelial cells during embryonic development at bulk and single-cell levels. Sci Signal 10, (2017).

125. M. Corada et al., Fine-Tuning of Sox17 and Canonical Wnt Coordinates the Permeability Properties of the Blood-Brain Barrier. Circ Res 124, 511–525 (2019).

126. R. N. Munji et al., Profiling the mouse brain endothelial transcriptome in health and disease models reveals a core blood-brain barrier dysfunction module. Nat Neurosci 22, 1892–1902 (2019).

127. N. Hannemann et al., The AP-1 Transcription Factor c-Jun Promotes Arthritis by Regulating Cyclooxygenase-2 and Arginase-1 Expression in Macrophages. J Immunol 198, 3605–3614 (2017).

128. Z. Han, D. L. Boyle, A. M. Manning, G. S. Firestein, AP-1 and NF-kappaB regulation in rheumatoid arthritis and murine collagen-induced arthritis. Autoimmunity 28, 197–208 (1998).

129. M. Guma, G. S. Firestein, c-Jun N-Terminal Kinase in Inflammation and Rheumatic Diseases. Open Rheumatol J 6, 220–231 (2012).

